# Role of stereochemistry on electron transport in peptides

**DOI:** 10.64898/2026.06.12.731974

**Authors:** Rajarshi Samajdar, Hemani Chhabra, Moeen Meigooni, Seungjoo Yi, Xiaolin Liu, Jason L. Wu, Emad Tajkhorshid, Charles M. Schroeder

**Affiliations:** Department of Chemical and Biomolecular Engineering, University of Illinois Urbana-Champaign, Urbana, IL 61801; Beckman Institute for Advanced Science and Technology, University of Illinois Urbana-Champaign, Urbana, IL 61801; Center for Biophysics and Quantitative Biology, University of Illinois Urbana-Champaign, Urbana, IL 61801; Department of Materials Science and Engineering, University of Illinois Urbana-Champaign, Urbana, IL 61801; Department of Chemical and Biological Engineering, Princeton University, Princeton, NJ 08544; Department of Chemistry, Princeton University, Princeton, NJ, 08544; Department of Chemistry, University of Illinois Urbana-Champaign, Urbana, IL 61801; Department of Bioengineering, University of Illinois Urbana-Champaign, Urbana, IL 61801

## Abstract

Stereochemistry underlies structure-function relationships across biology and materials science, ranging from proteins to electronic and spintronic materials. In this work, we investigate the electron transport properties of different oligopeptide stereoisomers using experiments and computational modeling. Single-molecule electronic experiments show that stereochemical modifications in tyrosine-based peptides lead to significant variations in molecular conductance along the peptide backbone due to enhanced stacking interactions and electronic coupling of aromatic side chains. In addition, stereochemical variations in alanine-based peptides give rise to changes in conductivity due to secondary structure interactions arising from β-turn conformations. All-atom molecular dynamics (MD) simulations and quantum mechanical calculations are used to understand the molecular origins of the effect of stereochemistry on the structural and electronic properties of peptides. Overall, this work shows that stereochemical modification of non-terminal amino acids effectively controls electron transport due to aromatic side chain interactions or secondary structure effects. These insights open new avenues for the molecular design of peptide-based electronic materials with enhanced function.

## Introduction

Stereochemistry governs the structure and function of synthetic and biological materials, influencing molecular conformation, intermolecular interactions, and the optical and electronic properties of a broad range of systems.^1,2^ Stereochemical effects play a central role in chemical reactions,^3^ transmission of photoexcited electrons through chiral media^4–7^, and electron transport through chiral materials under an applied bias.^8–13^ Prior work has focused on understanding the influence of stereochemistry on electron transport in biomolecules such as DNA^8,10^ and proteins,^14^ but systematic studies that directly connect stereochemical variation to sequence-dependent conformations and transport pathways in peptides remain limited. Despite recent progress, the relationship between stereochemistry, molecular sequence and structure, and electron transport in peptides is not fully understood.

Prior work on elucidating the effect of chirality on electron transport has focused on techniques such as magnetic conductive-probe atomic force microscopy (mCP-AFM)^10^, magnetoresistance in a film,^15^ and electrochemical measurements with a ferromagnetic electrode.^16^ These experiments are typically conducted at the bulk scale or through multiple molecular layers for complex biological systems. Additional work has focused on understanding molecular-scale electron transport in peptides, revealing that stereochemical variations in an α-helical peptide sequence of 22 amino acids alter the electronic properties using asymmetric Au and Ni metal electrodes.^17^ Nevertheless, the influence of stereochemistry on the electronic properties of short peptide sequences consisting of four to five amino acids (tetra- and pentapeptides) with symmetric metal electrodes has not been fully explored. Recent work showed that electron transport in peptides shows a two-state conductance profile due to the conformational flexibility of peptide backbones, with a high-conductance state arising due to a defined secondary structure (beta turn or 3_10_ helices) and a low-conductance state arising for extended peptide structures.^18^ From this perspective, investigating electron transport in short peptide sequences with different amino acid stereoisomers is critical for revealing how stereochemistry influences the electronic properties of peptides.

Bulk scale experiments have shown that stacking of aromatic amino acids such as tyrosine lead to enhancements in electronic conductivity.^19^ Building on these observations, we conjectured that stereochemical variations in peptide sequences could modulate molecular-scale electron transport in a conformation-dependent manner. In sequences containing consecutive tyrosine residues, replacing an internal L-tyrosine with D-tyrosine is predicted to yield structures with closer spatial alignment of adjacent aromatic side chains (**Scheme 1a**), strengthening electronic coupling in pi-stacking interactions and promoting a more defined electronic pathway along the peptide backbone. Conversely, in alanine-based sequences, replacing an internal L-alanine with D-alanine is thought to promote β-turn conformations in short peptides (**Scheme 1b**),^20,21^ thereby potentially enhancing electron transport through secondary structure-mediated pathways. In this way, we hypothesized that stereochemical modifications in tyrosine-based peptides would influence electron transport primarily along the molecular backbone, whereas analogous modifications in alanine-based sequences could modulate transport though intramolecular hydrogen-bonded pathways.

In this work, we investigate the role of stereochemistry on the electronic properties of short peptides using a combination of single molecule experiments, bulk scale spectroscopic characterization, molecular dynamics (MD) simulations, and quantum mechanical (QM) calculations. A scanning tunneling microscope-break junction (STM-BJ) technique^18,22–26^ is used to experimentally characterize the molecular charge transport behavior of oligopeptides defined by different non-terminal amino acid chiral isomers. Results from single-molecule electronics experiments show that stereochemical changes consisting of alternating L- and D-residues in tyrosine and alanine-based pentapeptides enhance molecular conductance relative to the homochiral analogs. All-atom MD simulations demonstrate that heterochiral peptides, in which non-terminal amino acids have alternating chirality, exhibit enhanced pi-stacking interactions between the aromatic side chains of tyrosine residues. On the other hand, heterochiral pentapeptides based on alanine display pronounced secondary structure interactions due to a favorable β-turn conformation. QM calculations are further used to validate experimental results for molecular-scale electron transport. Results from single-molecule experiments on tetrapeptides show a similar yet more subtle role of stereochemical modifications on conductivity of tetrapeptides. Overall, our work illustrates that changes in stereochemistry directly affect electron transport in peptides via primary or secondary structure-mediated pathways.

**Scheme 1:**
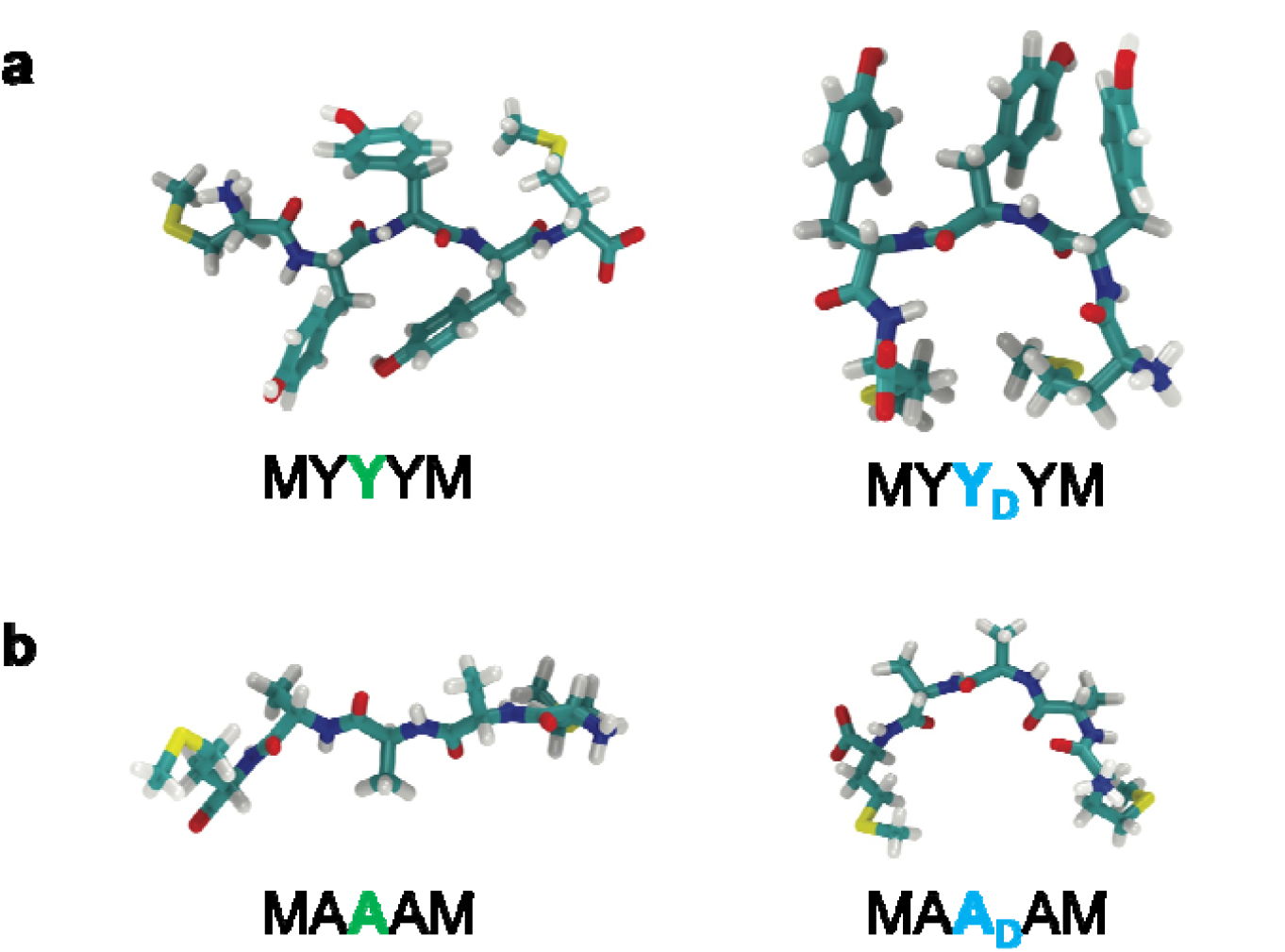
Understanding the role of stereochemistry on electron transport in peptides. (a) Substituting internal L-tyrosine with D-tyrosine residues is predicted to yield a closer spatial alignment of aromatic side chains, providing a defined electronic pathway along the peptide backbone. (b) Substituting internal L-alanine with D-alanine is predicted to favor a β-turn conformations,^20,21^ leading to a more pronounced electronic pathway due to secondary structure interactions. M, Y, and A denote the L-amino acids methionine, tyrosine, and alanine, respectively, whereas Y_D_ and A_D_ denote the corresponding D-amino acids.

## Results and Discussion

### Molecular design and characterization

Tetra- and pentapeptides were designed with non-polar aliphatic groups and aromatic R groups, featuring variations in the stereochemistry of the chiral centers on the alpha-carbons of non-terminal amino acids (**Figures 1a,b** and **Supplementary Figures 1-10)**. The chemical library consists of homochiral peptides (non-terminal amino acids with identical stereochemistry) and heterochiral peptides (non-terminal amino acids with alternating stereochemistry) based on tyrosine (**Figure 1a**) and alanine (**Figure 1b**). We began by characterizing peptides using UV-visible and fluorescence spectroscopy. Tyrosine-based peptides exhibit a distinct absorption peak at 275 nm with shoulders at 282 nm in the UV-visible spectrum (**Supplementary Figure 11**), along with an emission peak near 305 nm in the fluorescence spectrum (**Supplementary Figure 12)**, consistent with the characteristic optical features of aromatic residues for peptides reported in prior literature.^27,28^ In contrast, alanine-based peptides exhibit no distinct peaks in the aromatic region (**Supplementary Figures 13**,**14**) with an absorption around 190-220 nm due to the C=O transitions of the peptide backbone.^29,30^

**Figure 1:**
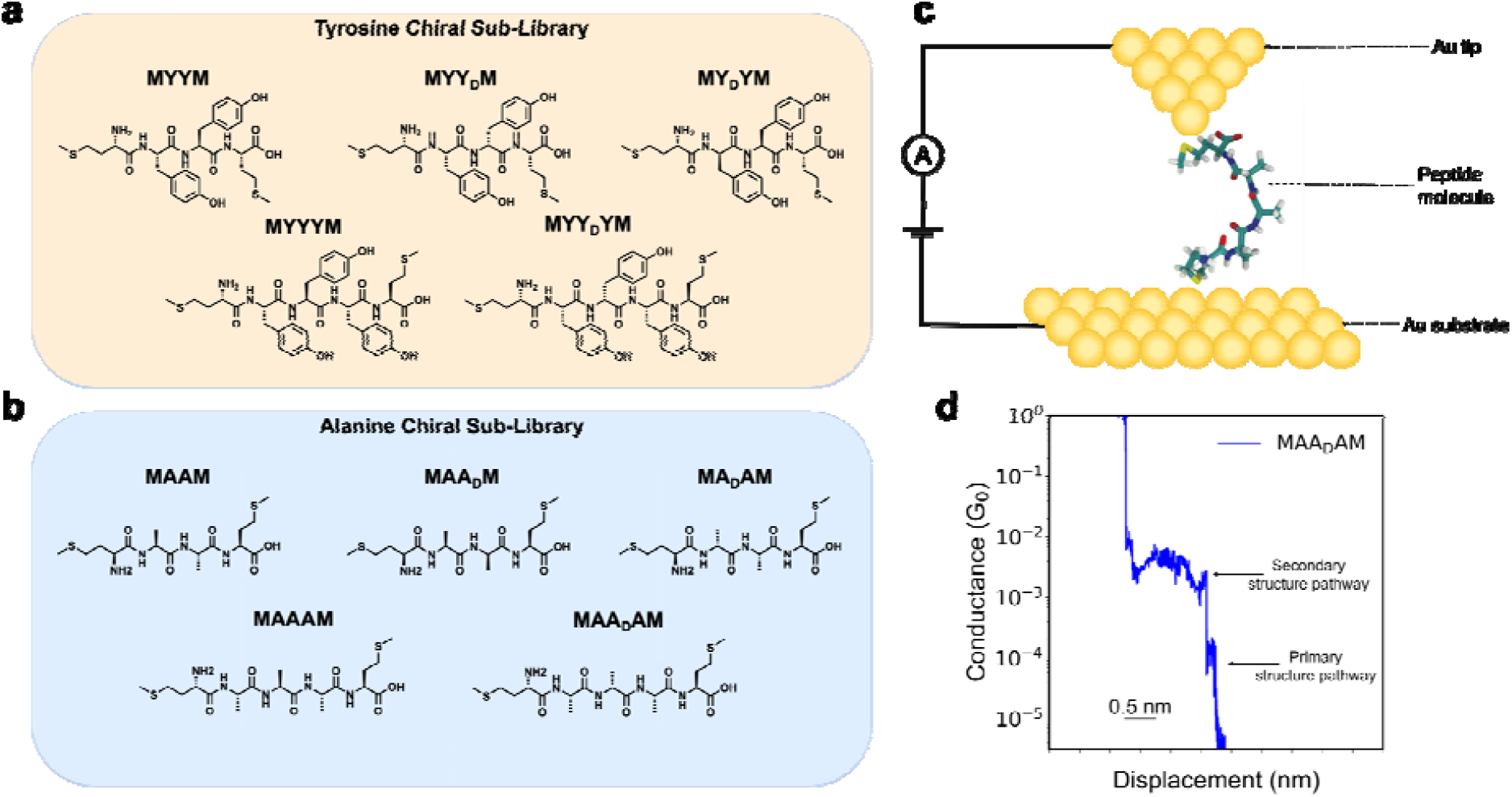
Molecular library of chiral peptides studied in this work. (a) Tyrosine-based chiral peptide sub-library. (b) Alanine-based chiral peptide sub-library. (c) Schematic of scanning tunneling microscope-break junction (STM-BJ) experiment with the peptide MAA_D_AM. (d) Characteristic single-molecule conductance trace for peptide MAA_D_AM, indicating a high-conductance pathway and a low-conductance pathway arising from secondary and primary structures, respectively.^18^ M, Y, and A denote the L-amino acids methionine, tyrosine, and alanine, respectively, whereas Y_D_ and A_D_ denote the corresponding D-amino acids.

To further understand the role of stereochemistry on molecular conformations, we performed two-dimensional (2D) nuclear magnetic resonance (NMR) spectroscopy. Diffusion-ordered spectroscopy (DOSY) NMR is used to determine diffusion coefficients, providing information on molecular size and shape in solution.^31,32^ We performed DOSY NMR experiments on tyrosine-based peptide sequences, as differences in these peptides could arise due to interactions between aromatic side chains (**Scheme 1a**). Results from DOSY NMR experiments on peptides MYYYM and MYY_D_YM revealed similar diffusion coefficients, indicating comparable molecular sizes and shapes in solution (**Supplementary Figure 15**,**16**). Nuclear Overhauser effect spectroscopy (NOESY) NMR was also used to probe through-space interactions between protons, revealing local conformational differences induced by stereochemistry.^31,33^ NOESY NMR were performed on alanine-based peptide sequences, as differences in tertiary conformation could arise due to distinct folded structures of these peptide isomers (**Scheme 1b**). NOESY NMR results for MAAAM and MAA_D_AM showed no observable cross-peaks (**Supplementary Figures 17**), suggesting the absence of close proton–proton spatial contacts in solution phase. However, methods such as NMR reflect ensemble-averaged conformations over time and across the populations of molecules in solution, and may not capture transient or folded states, if they exist in only minor sub-populations. To further probe the presence of folded conformations, we performed circular dichroism (CD) spectroscopy,^18,34^ which is more sensitive to secondary structure and can detect subtle changes in backbone conformation induced by chirality.^35,36^

Circular dichroism (CD) spectra (**Supplementary Figures 18-21)** indicate the presence of hydrogen bonding (H-bonding) interactions for the tetra- and pentapeptides, showing spectral features expected for beta turn or 3_10_ helices, such as a maximum or minimum around 200-210 nm and a shoulder or small peak around 220 nm.^37,38^ CD spectral features for 3_10_ helices are qualitatively different than the spectral features observed for alpha helices, beta sheets, or random coils.^39^ Additionally, pronounced differences are observed between homochiral and heterochiral peptide sequences. Several heterochiral peptides, including MYY_D_YM, MA_D_AM, and MAA_D_AM, display spectral features in the 190-200 nm region (**Supplementary Figure 19-21**) that differ from their homochiral counterparts. Furthermore, the CD spectrum of MY_D_YM appears as a shifted mirror image of that of MYYM (**Supplementary Figure 18**), highlighting the role of amino acid stereochemistry on peptide conformation. These observations indicate that peptides composed of the same amino acid sequence but differing in the stereochemistry of non-terminal residues adopt markedly different conformational ensembles. Such differences likely arise from changes in side-chain orientation, intramolecular interactions, and three-dimensional conformational arrangements induced by stereochemical variation. We next performed single-molecule measurements to understand how peptide structural variations affect electron transport.

### Single-molecule conductance measurements

Tetra- and pentapeptides were designed with a methionine residue at the N- and C-termini. Methionine contains a methyl sulfide (-S-CH_3_) group that readily binds to gold,^40^ thereby providing electrical contacts to metal electrodes in the STM-BJ setup (**Figure 1c**). All STM-BJ measurements on peptides were carried out in water (peptide concentration <1 mM). Prior work^18^ revealed a two-state molecular conductance behavior of homochiral peptides due to the conformational flexibility of peptide backbones, with a high-conductance state arising due to secondary structure interactions (beta turn or 3_10_ helices) and a low-conductance state occurring for extended peptide structures (**Figure 1d**). The present work aims to understand how changes in stereochemistry affect the molecular electronic properties of peptides.

We began by characterizing the electron transport behavior of tyrosine-based tetrapeptides. Our results show that the high conductance state of tyrosine-based homochiral tetrapeptides (MYYM) is similar to the high conductance state of heterochiral tetrapeptides (MYY_D_M and MY_D_YM) around ∼10^−2.8^ *G*/*G*_*0*_, suggesting that the electron transport pathway mediated by secondary structure interactions is not significantly altered by stereochemical changes in tyrosine-based sequences (**Supplementary Figure 22** and **Supplementary Table S1**). In contrast, the low conductance state, corresponding to the electron transport pathway along the primary peptide backbone, shows significant variation between chiral isomers in tyrosine-based peptides, with the heterochiral sequence MYY_D_M exhibiting a conductance value ∼10^0.5^ *G*/*G*_*0*_ smaller than the homochiral peptide MYYM (**Supplementary Figure 22** and **Supplementary Table S1**).

The effect of stereochemistry on molecular conductance was found to be more pronounced in tyrosine-based pentapeptides (**Figure 2**). Characteristic single-molecule traces (**Figure 2a**) show that stereochemical changes in tyrosine-based pentapeptides result in a pronounced low conductance feature. One-dimensional and two-dimensional molecular conductance histograms generated for ensembles of >5000 single molecules (**Figures 2b,c,d**) reveal that the low conductance state is affected by stereochemical changes in tyrosine, with the peak molecular conductance value for MYY_D_YM around ∼10^−4.8^ *G*/*G*_*0*_ compared to the peak molecular conductance of MYYYM around ∼10^−4.0^ *G*/*G*_*0*_ (**Supplementary Table S2**). Notably, this decrease in conductance is accompanied by an increase in the molecular junction separation distance for MYY_D_YM, suggesting that stereochemical modification alters both the pathway length and electronic coupling along the peptide backbone. Based on these results, we posit that alternating stereochemistry of internal amino acid residues in tyrosine-based sequences results in enhanced spatial alignment of aromatic side chains, thereby promoting pi-stacking interactions, enhancing electronic coupling, and creating a defined electron transport pathway along the primary backbone. This hypothesis is evaluated using molecular simulations and modeling, as discussed below.

**Figure 2:**
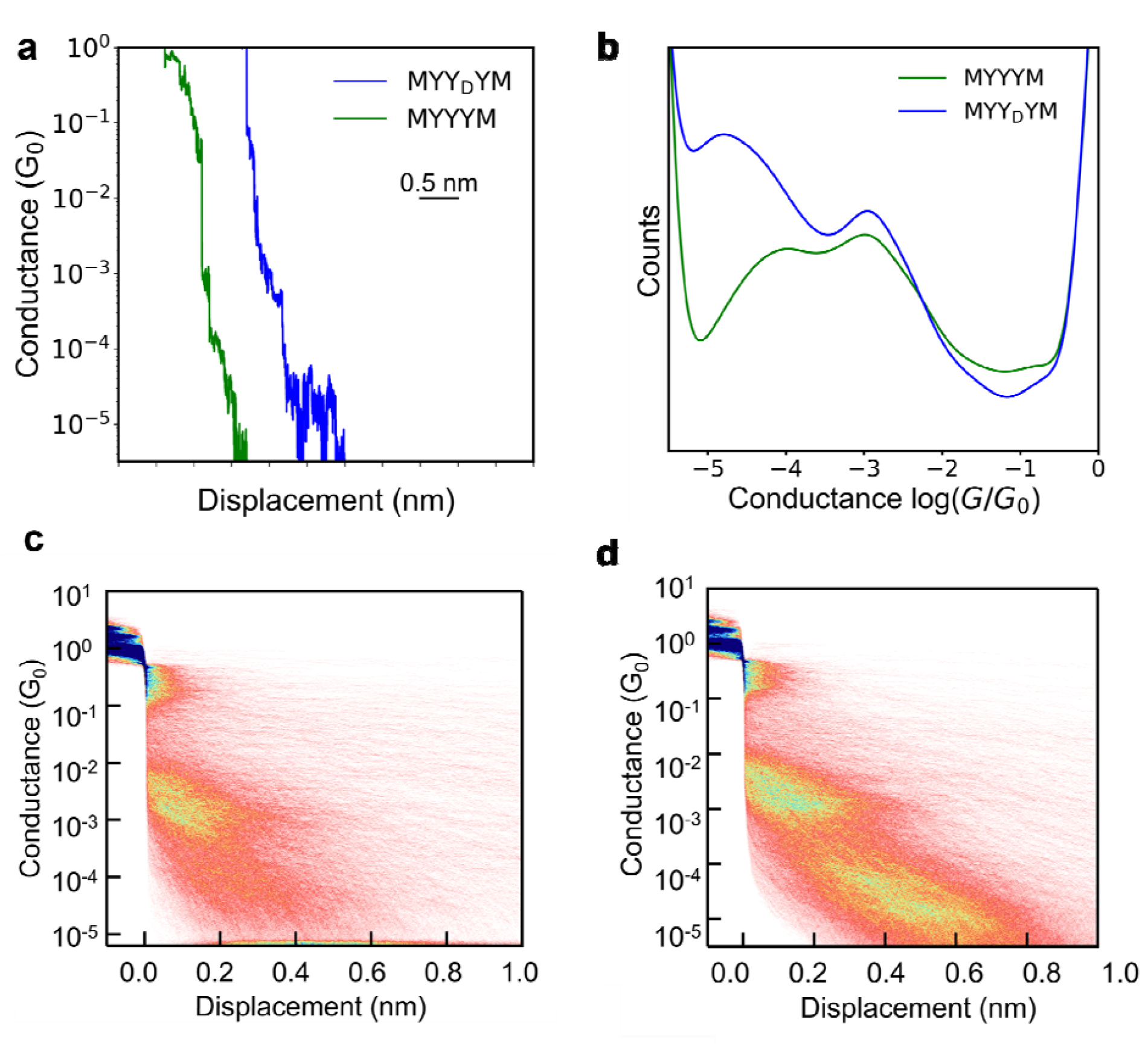
Single molecule electronic measurements for tyrosine-based pentapeptides. (a) Characteristic single molecule traces for MYYYM and MYY_D_YM. (b) 1D conductance histograms for homochiral peptide MYYYM and heterochiral peptide MYY_D_YM. 2D conductance histograms for (c) MYYYM and (d) MYY_D_YM. M, Y, and A denote the L-amino acids methionine, tyrosine, and alanine, respectively, whereas Y_D_ and A_D_ denote the corresponding D-amino acids. All data were obtained using 0.1 mM concentrations of peptoids in water at 250 mV applied bias across ensembles of at least 5000 single molecules.

We next performed single-molecule electronic experiments on alanine-based tetra- and pentapeptides. Our results show that alanine-based homochiral and heterochiral tetrapeptides show similar high and low conductance features (**Supplementary Figure 23** and **Supplementary Table S3**). However, alanine-based heterochiral tetrapeptide sequences show a higher occurrence of the high conductance state compared to homochiral alanine-containing tetrapeptides (**Supplementary Figure 23a**). Characteristic single-molecule traces for alanine-based pentapeptides (**Figure 3a**) show an enhanced high-conductance feature for heterochiral peptides with alternating stereochemistry of internal amino acid residues, which likely arises due to enhanced secondary structure interactions. One-dimensional and two-dimensional molecular conductance histograms generated across ensembles of >5000 single molecules (**Figures 3b,c,d** and **Supplementary Table S4**) reveal higher counts and a longer molecular junction distance for MAA_D_AM compared to MAAAM. Alternating the chirality of alanine residues in a peptide sequence is known to promote favorable β-turn conformations,^20,21^ which could support an enhanced electronic pathway due to secondary structure interactions. To further understand these findings, computational modeling is used to rationalize the experimental results.

**Figure 3:**
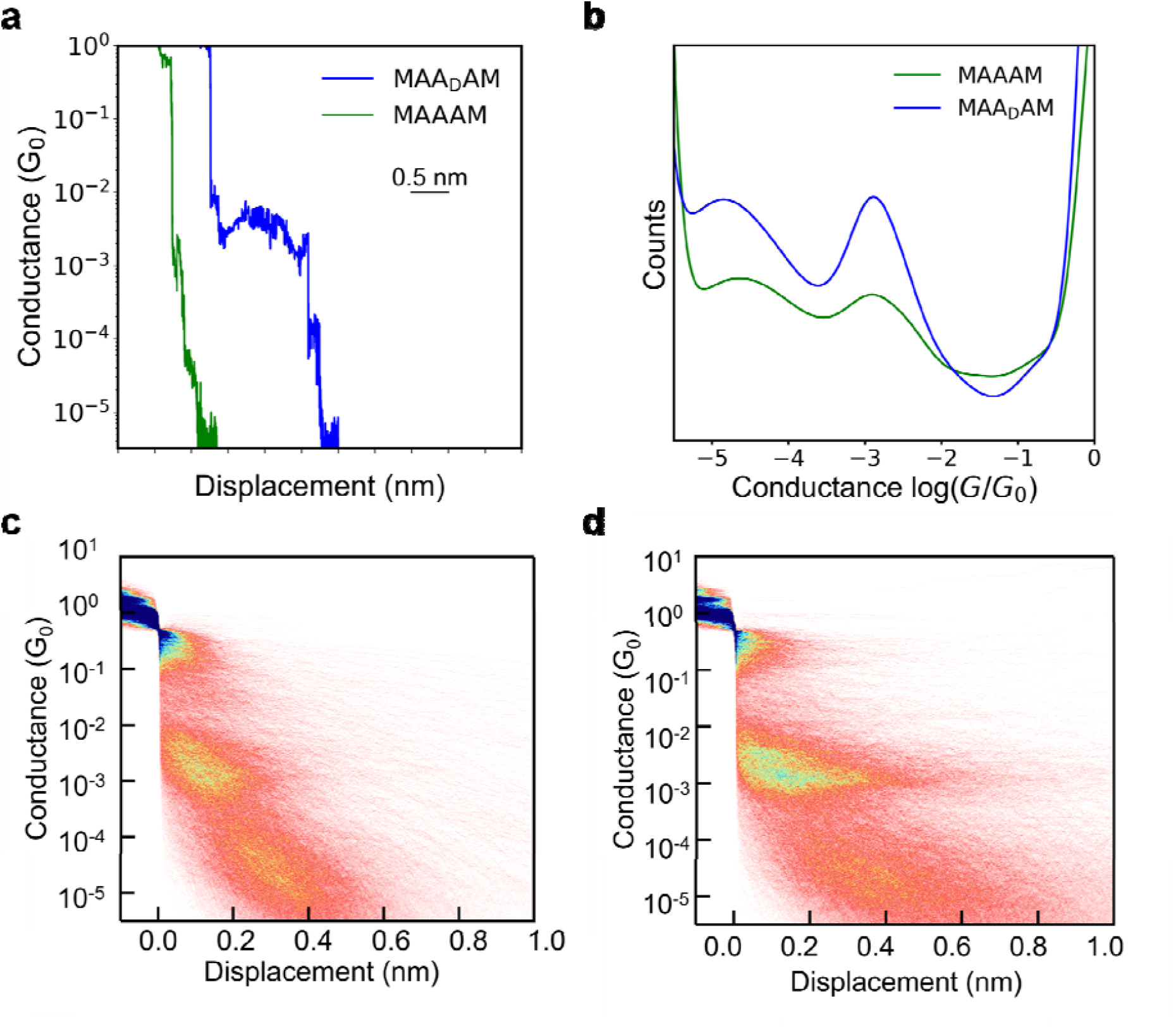
Single molecule electronic measurements for alanine-based pentapeptides. (a)Characteristic single molecule traces for MAAAM and MAA_D_AM. (b) 1D conductance histogram for MAAAM and MAA_D_AM. 2D conductance histogram for (c) MAAAM and (d) MAA_D_AM. M, Y, and A denote the L-amino acids methionine, tyrosine, and alanine, respectively, whereas Y_D_ and A_D_ denote the corresponding D-amino acids. All data were obtained using 0.1 mM concentrations of peptoids in water at 250 mV applied bias across ensembles of at least 5000 single molecules.

### Molecular dynamics (MD) simulations

To understand the role of molecular conformation on electron transport, we performed MD simulations for tyrosine and alanine-based peptides (**Figure 4a**). MD simulations were performed in explicit solvent with a series of custom potentials to implicitly represent interactions between peptides and gold electrodes, as described in prior work^18^ (**Supplementary Figure 24**). The projection of the end-to-end distance (sulfur anchor-to-anchor distance on terminal methionines) of the peptide along the experimental pulling axis was harmonically restrained to a series of distances (6 Å, 9 Å, and 12 Å), allowing the peptide to adopt an ensemble of conformations. MD simulations for homochiral peptides, MAAAM and MYYYM, show that 2 → 5 backbone hydrogen bonds (H-bonds) form with remarkable consistency during the 6 Å end-to-end holding distance, but the interactions vanish when the end-to-end distance is 12 Å (**Supplementary Figure 25**). This observation is consistent with our experimental findings in this work and prior work based on single-molecule experiments.^18^ We also analyzed MD simulation results to elucidate the origin of the enhanced low-conductance state in MYY_D_YM compared to MYYYM, presumably associated with electron transport along the primary backbone, and the origin of the pronounced high-conductance state in MAA_D_AM compared to MAAAM, presumably associated with electron transport due to secondary structure interactions.

**Figure 4:**
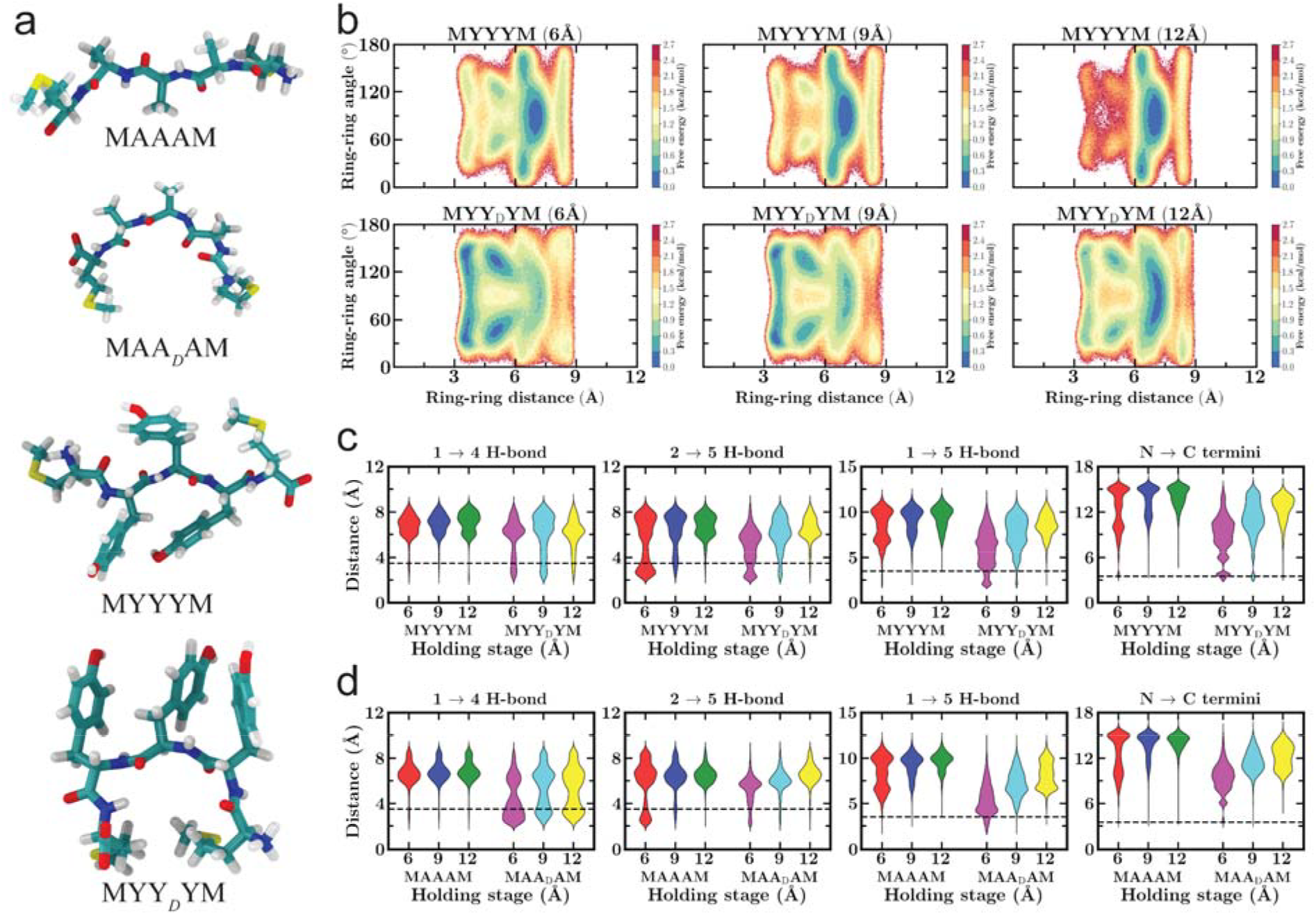
MD simulation results for homochiral and heterochiral pentapeptides. (a) Snapshots for alanine and tyrosine-based pentapeptides studied in this work. (b) Free energy plots for tyrosine-based pentapeptides as a function of the distance and angle between the aromatic side chains, indicating increased interactions between phenyl rings in heterochiral peptides as compared to homochiral peptides. (c), (d) Violin plots showing backbone H-bonding distance distribution for tyrosine and alanine based pentapeptides. M, Y, and A denote the L-amino acids methionine, tyrosine, and alanine, respectively, whereas Y_D_ and A_D_ denote the corresponding D-amino acids. A black dotted line in the violin plots indicate the cutoff for intermolecular H-bonding.

The energy profiles for tyrosine-based homochiral and heterochiral sequences as a function of the distance and angle between the aromatic side chains indicate closer spatial alignment and enhanced interactions between the phenyl rings in MYY_D_YM compared to MYYYM (**Figure 4b**). Violin plots of backbone H-bonding distances for tyrosine-based peptides indicate a more pronounced population at larger end-to-end distances (9 Å and 12 Å) for MYY_D_YM compared to MYYYM (**Figure 4c**). These results indicate that enhanced aromatic interactions between the tyrosine side chains result in a more dominant electron transport pathway along the primary structure, consistent with results from single molecule experiments. Overall, MD simulations and analyses based on backbone H-bonding, as well as free energy landscapes describing the orientation and alignment of phenyl side chains, support the molecular and structural basis for the enhanced electronic pathway along the primary structure observed in heterochiral tyrosine-based peptides.

Violin plots for MAAAM and MAA_D_AM based on backbone H-bonding are shown in **Figure 4d**. Our results reveal that MAA_D_AM has a prominent population at small end-to-end distances (6 Å) compared to MAAAM due to a 1 → 4 H-bonding interaction, indicating increased folding (β-turn or 3_10_ helix character), whereas MAAAM predominantly exhibits a 2 → 5 H-bonding interaction (**Figure 4d**). From this perspective, these results suggest that the probability of forming secondary structures, such as 3_10_ helices, strongly influences electronic functionality depending on the conformation adopted by the peptide. For instance, a 1→4 H-bonding interaction can differ from a 2→5 H-bonding interaction. Overall, results from all-atom MD simulations suggest that the heterochiral peptide MAA_D_AM has a prominent electron transport pathway due to secondary structure interactions owing to more favorable β-turn or 3_10_ helical formations compared to the homochiral peptides, consistent with single molecule experimental results. Next, quantum mechanical calculations and non-equilibrium Green’s function–density functional theory (NEGF-DFT) simulations were used to further to rationalize experimental results.

### NEGF-DFT and quantum calculations

To complement the single-molecule conductance measurements and MD simulations, density functional theory (DFT) calculations were performed to understand the role of stereochemistry on the electronic structure of tyrosine- and alanine-based peptides. These calculations were carried out on representative peptide conformations obtained from the MD simulations, corresponding to the low-conductance state of the tyrosine-based peptides and the high-conductance state of the alanine-based peptides observed in the STM-BJ experiments. Results from frontier molecular orbital analysis (**Supplementary Figures 26-29**) show that stereochemical alterations do not significantly affect the energy alignment of the HOMO-1, HOMO, LUMO, and LUMO+1 levels within each peptide pair (MAAAM vs MAADAM and MYYYM vs MYYDYM), as shown in **Supplementary Table S5**. Based on these observations, we propose that differences in electron transport between sequences of differing stereochemistry are governed by conformational variations, supported by the MD simulations, rather than significant differences in frontier orbital energies.

To further understand the role of amino acid stereochemistry on the electronic properties of peptides, we performed projected density of states (PDOS) calculations (**Supplementary Figures 30-31**). PDOS analysis demonstrates that stereochemical inversion redistributes orbital character near the frontier states, as evidenced by changes in the relative contributions of different atomic sites without significant shifts in energy. Overall, results from DFT modeling highlight the importance of structural and conformational effects, rather than significant changes in electronic level alignment, in governing electron transport through these peptides.

We next performed electron transmission calculations on tyrosine and alanine-based peptides. Representative molecular structures corresponding to the most probable conformations generated by MD were used in non-equilibrium Green’s function-density functional theory (NEGF-DFT) calculations to enable direct comparison between theory and experimental results. Electron transmission calculations performed on representative peptide conformations corresponding to the low-conductance state reveal a nearly fourfold difference in transmission at the Fermi energy between MYYYM and MYY_D_YM (**Supplementary Figures 32a**,**b**), consistent with the trends observed in single-molecule break-junction experiments (**Supplementary Table 6**). Electron transmission calculations for the alanine-based peptides MAAAM and MAA_D_AM, corresponding to the high-conductance state, reveal comparable transmission values at the Fermi energy (**Supplementary Figures 32c**,**d**), consistent with the experimental observations (**Supplementary Table S7**). Taken together, results from NEGF-DFT and quantum calculations show that alternating stereochemical sequences of tyrosine and alanine-based amino acids modulate electron transport through conformationally dependent changes in electronic coupling, rather than through shifts in frontier orbital energies.

## Conclusions

In this work, the electronic properties of oligopeptides defined by different non-terminal amino acid stereoisomers are characterized using experiments and modeling. Results from single-molecule experiments and computational modeling reveal that stereochemical modifications in tyrosine-based sequences lead to significant variations in electronic conductivity along the primary structure due to increased interactions among the aromatic side chains of tyrosine residues. Moreover, chirality changes in alanine-based pentapeptides result in significant variations in electronic conductivity along the secondary structure due to more favorable β-turn conformations. All-atom MD simulations are used to analyze backbone H-bonding interactions and free energy landscapes describing the orientation and alignment of phenyl side chains, thereby providing a rational basis for understanding the putative pathways for electron transport observed in the experimental results. Quantum mechanical calculations, including NEGF-DFT, further establish a direct connection between structure, electronic structure, and transport behavior.

Overall, our work demonstrates the tuning of electronic properties at the molecular scale by varying the stereochemistry of specific amino acids within a peptide backbone, leading to enhanced electron transport due to primary structure or secondary structure interactions. The insights gained from this work provide a framework for understanding how stereochemistry influences electron transport in biomolecular systems and offer design principles for engineering longer peptide and protein-inspired sequences with tailored electronic properties. By strategically varying the stereochemistry of individual amino acids, it may be possible to modulate peptide conformation, self-assembly behavior, intermolecular interactions, and charge-transport pathways, thereby enabling systematic control over electronic currents and functionality in bioelectronic materials. Moving forward, extending this approach to longer and more complex sequences, incorporating environmental effects such as solvent^26^ will be essential to fully elucidate the interplay between structure and electronic function. Broadly, these insights could be used to open new avenues for the design of bio-inspired electronic materials and molecular devices, where stereochemistry can be leveraged to engineer tunable and directional charge transport pathways.

## Acknowledgements

This work was supported by the U.S. Department of Energy, Office of Science, Basic Energy Sciences under Award No. DE-SC0022035 for X.L., M.M., J.S.M., E.T., and C.M.S. and the Army Research Office under Cooperative Agreement Number W911NF-22-2-0246 for R.S. and C.M.S. E.T. also acknowledges support from the National Institutes of Health, grant R24-GM145965. The views and conclusions contained in this document are those of the authors and should not be interpreted as representing the official policies, either expressed or implied, of the Army Research Office or the U.S. Government. The U.S. Government is authorized to reproduce and distribute reprints for Government purposes notwithstanding any copyright notation herein. This work was supported by the Molecule Maker Lab Institute, an AI Research Institutes program supported by the US National Science Foundation under grant no. 2019897 for S.Y. H.C. acknowledges support from the Beckman Fellowship, University of Illinois Urbana Champaign. We thank Jeffrey S. Moore for useful discussions.

## Author contributions

R.S. and C.M.S. conceived this study. R.S. designed the peptide sequences, performed the STM-BJ experiments and data analysis. M.M. and H.C. performed the MD simulations and analysis. S.Y. performed NMR characterization. X.L. and R.S. performed bulk scale spectroscopy and circular dichroism experiments. R.S and J.W performed the NEGF-DFT and quantum mechanical calculations. The manuscript was written by R.S. and C.M.S. with contribution from all authors.

## Competing Interests

The authors declare no competing interests.

## Additional Information

Supplementary information contains methods, mass spectrometry data, UV-visible and fluorescence spectroscopy data, two-dimensional (2D) nuclear magnetic resonance (NMR) spectroscopy, circular dichroism (CD) spectroscopy data, quantum mechanical calculations, and additional molecular scale experimental results.

## Table of Contents Graphic

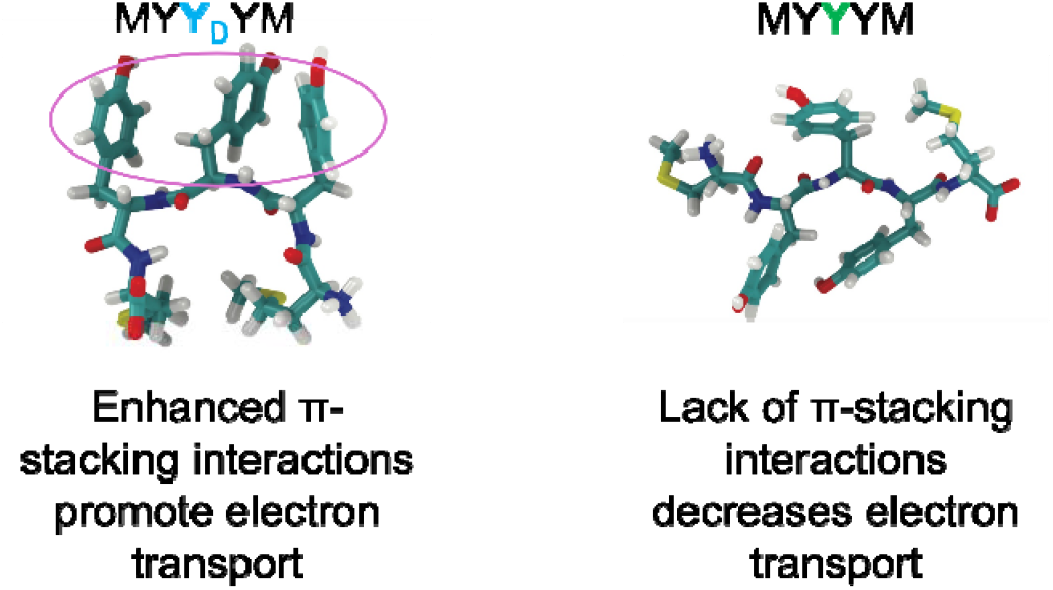

## Notes

### Competing Interest Statement

The authors have declared no competing interest.

## References

1. Barron, L. D. Symmetry and chirality: where physics shakes hands with chemistry and biology. Israel Journal of Chemistry 2021, 61 (9-10), 517–529.

2. Evers, F.; Aharony, A.; Bar-Gill, N.; Entin-Wohlman, O.; Hedegård, P.; Hod, O.; Jelinek, P.; Kamieniarz, G.; Lemeshko, M.; Michaeli, K.; Mujica, V.; Naaman, R.; Paltiel, Y.; Refaely-Abramson, S.; Tal, O.; Thijssen, J.; Thoss, M.; van Ruitenbeek, J. M.; Venkataraman, L.; Waldeck, D. H.; Yan, B.; Kronik, L. Theory of chirality-induced spin selectivity: progress and challenges. Advanced Materials 2022, 34 (13), 2106629.

3. Kumar, A.; Capua, E.; Kesharwani, M. K.; Martin, J. M.; Sitbon, E.; Waldeck, D. H.; Naaman, R. Chirality-induced spin polarization places symmetry constraints on biomolecular interactions. Proceedings of the National Academy of Sciences 2017, 114 (10), 2474–2478.

4. Ray, K.; Ananthavel, S. P.; Waldeck, D. H.; Naaman, R. Asymmetric scattering of polarized electrons by organized organic films of chiral molecules. Science 1999, 283 (5403), 814–816.

5. Zhao, B. S.; Meijer, G.; Schöllkopf, W. Quantum reflection of He2 several nanometers above a grating surface. Science 2011, 331 (6019), 892–894.

6. Carmeli, I.; Skakalova, V.; Naaman, R.; Vager, Z. Magnetization of chiral monolayers of polypeptide: a possible source of magnetism in some biological membranes. Angewandte Chemie International Edition 2002, 114 (5), 787–790.

7. Niño, M. Á.; Kowalik, I. A.; Luque, F. J.; Arvanitis, D.; Miranda, R.; de Miguel, J. J. Enantiospecific spin polarization of electrons photoemitted through layers of homochiral organic molecules. Advanced Materials 2014, 26 (44), 7474–7479.

8. Abendroth, J. M.; Nakatsuka, N.; Ye, M.; Kim, D.; Fullerton, E. E.; Andrews, A. M.; Weiss, P. S. Analyzing spin selectivity in DNA-mediated charge transfer via fluorescence microscopy. ACS Nano 2017, 11 (7), 7516–7526.

9. Lu, H.; Xiao, C.; Song, R.; Li, T.; Maughan, A. E.; Levin, A.; Brunecky, R.; Berry, J. J.; Mitzi, D. B.; Blum, V.; Beard, M. C. Highly distorted chiral two-dimensional tin iodide perovskites for spin polarized charge transport. Journal of the American Chemical Society 2020, 142 (30), 13030–13040.

10. Xie, Z.; Markus, T. Z.; Cohen, S. R.; Vager, Z.; Gutierrez, R.; Naaman, R. Spin specific electron conduction through DNA oligomers. Nano Letters 2011, 11 (11), 4652–4655.

11. Alam, K. M.; Pramanik, S. Spin filtering with poly-T wrapped single wall carbon nanotubes. Nanoscale 2017, 9 (16), 5155–5163.

12. Kulkarni, C.; Mondal, A. K.; Das, T. K.; Grinbom, G.; Tassinari, F.; Mabesoone, M.F. J.; Meijer, E. W.; Naaman, R. Highly efficient and tunable filtering of electrons’ spin by supramolecular chirality of nanofiber-based materials. Advanced Materials 2020, 32 (7), 1904965.

13. Alam, K. M.; Pramanik, S. Spin filtering through single-wall carbon nanotubes functionalized with single-stranded DNA. Advanced Functional Materials 2015, 25 (21), 3210–3218.

14. Gupta, R.; Chinnasamy, H. V.; Sahu, D.; Matheshwaran, S.; Sow, C.; Mondal, P. C. Spin-dependent electrified protein interfaces for probing the CISS effect. The Journal of Chemical Physics 2023, 159 (2), 024101.

15. Ben Dor, O.; Morali, N.; Yochelis, S.; Baczewski, L. T.; Paltiel, Y. Local light-induced magnetization using nanodots and chiral molecules. Nano Letters 2014, 14 (11), 6042–6049.

16. Mishra, D.; Markus, T. Z.; Naaman, R.; Kettner, M.; Göhler, B.; Zacharias, H.; Friedman, N.; Sheves, M.; Fontanesi, C. Spin-dependent electron transmission through bacteriorhodopsin embedded in purple membrane. Proceedings of the National Academy of Sciences 2013, 110 (37), 14872–14876.

17. Aragonès, A. C.; Medina, E.; Ferrer-Huerta, M.; Gimeno, N.; Teixidó, M.; Palma, J. L.; Tao, N.; Ugalde, J. M.; Giralt, E.; Díez-Pérez, I.; Mujica, V. Measuring the spin-polarization power of a single chiral molecule. Small 2017, 13 (2), 1602519.

18. Samajdar, R.; Meigooni, M.; Yang, H.; Li, J.; Liu, X.; Jackson, N. E.; Mosquera, M. A.; Tajkhorshid, E.; Schroeder, C. M. Secondary structure determines electron transport in peptides. Proceedings of the National Academy of Sciences 2024, 121 (32), e2403324121.

19. Shipps, C.; Kelly, H. R.; Dahl, P. J.; Yi, S. M.; Vu, D.; Boyer, D.; Glynn, C.; Sawaya, M. R.; Eisenberg, D.; Batista, V. S.; Malvankar, N. S. Intrinsic electronic conductivity of individual atomically resolved amyloid crystals reveals micrometer-long hole hopping via tyrosines. Proceedings of the National Academy of Sciences 2021, 118 (2), e2014139118.

20. Yan, Y.; Erickson, B. W.; Tropsha, A. Free energies for folding and refolding of four types of β turns: simulation of the role of D/L chirality. Journal of the American Chemical Society 1995, 117 (29), 7592–7599.

21. Yan, Y.; Tropsha, A.; Hermans, J.; Erickson, B. W. Free energies for refolding of the common β turn into the inverse-common β turn: simulation of the role of D/L chirality. Proceedings of the National Academy of Sciences 1993, 90 (16), 7898–7902.

22. Liu, X.; Yang, H.; Harb, H.; Samajdar, R.; Woods, T. J.; Lin, O.; Chen, Q.; Romo, A.B.; Rodríguez-López, J.; Assary, R. S.; Moore, J. S.; Schroeder, C. M. Shape-persistent ladder molecules exhibit nanogap-independent conductance in single-molecule junctions. Nature Chemistry 2024, 16, 1–9.

23. Samajdar, R.; Yang, H.; Yi, S.; Wang, C.-I.; Putnam, S. T.; Pence, M. A.; Lindsay, G. S.; Meigooni, M.; Liu, X.; Ren, J.; Moore, J. S.; Tajkhorshid, E.; Gewirth, A. A.; Rodríguez-López, J.; Jackson, N. E.; Schroeder, C. M. Electrochemically mediated Au–C(sp2) anchors for molecular electronics. The Journal of Physical Chemistry C 2025, 129 (39), 17458–17471.

24. Prempin, B.; Rajarshi Samajdar; Chhabra H.; Meigooni, M.; Aksimentiev, A.; Tajkhorshid, E.; Moore, J. S.; Schroeder, C. M. Single-molecule electron transport in peptoids. The Journal of Physical Chemistry B 2026, 130 (11), 3054–3064.

25. Samajdar, R.; Liu, X.; Kuyama, K.; Kidokoro, Y.; Takeda, F.; Okamoto, I.; Kawahata, M.; Katagiri, K.; Moore, J. S.; Tanatani, A.; Schroeder, C. M. Aromatic amide foldamers show conformation-dependent electronic properties. ChemPhysChem 2025, 26 (24), e202500672.

26. Samajdar, R.; Nadeem, H.; Moghe, N.; Shukla, D.; & Schroeder, C. M. (2026). Solvent Environment Influences Molecular Conformation and Electron Transport in Peptides. The Journal of Physical Chemistry Letters 202617(21), 6004–6013.

27. Antosiewicz, J. M.; Shugar, D. UV–Vis spectroscopy of tyrosine side groups in studies of protein structure. Part 2: Selected applications. Biophysical Reviews 2016, 8, 163–177.

28. Lucas, L. H.; Ersoy, B. A.; Kueltzo, L. A.; Joshi, S. B.; Brandau, D. T.; Thyagarajapuram, N.; Peek, L. J.; Middaugh, C. R. Probing protein structure and dynamics by second-derivative ultraviolet absorption analysis of cation–π interactions. Protein Science 2006, 15, 2228–2243.

29. Sharma, B.; Asher, S. A. UV resonance Raman investigation of the conformations and lowest energy allowed electronic excited states of tri- and tetraalanine: Charge transfer transitions. The Journal of Physical Chemistry B 2010, 114, 6661–6668.

30. Heller, A.; Rönitz, O.; Barkleit, A.; Bernhard, G.; Ackermann, J.-U. Complexation of europium (III) with the zwitterionic form of amino acids studied with ultraviolet–visible and time-resolved laser-induced fluorescence spectroscopy. Applied Spectroscopy 2010, 64, 930–935.

31. Doyen, C.; Larquet, E.; Coureux, P.-D.; Frances, O.; Herman, F.; Sablé, S.; Burnouf, J.-P.; Sizun, C.; Lescop, E. Nuclear magnetic resonance spectroscopy: A multifaceted toolbox to probe structure, dynamics, interactions, and real-time in situ release kinetics in peptide-liposome formulations. Molecular Pharmaceutics 2021, 18, 2521–2539.

32. Wang, K.; Chen, K. Direct assessment of oligomerization of chemically modified peptides and proteins in formulations using DLS and DOSY-NMR. Pharmaceutical Research 2023, 40, 1329–1339.

33. Filia, S.; Tambunan, U. S. F.; Kuczera, K.; Siahaan, T. J. Structure of a cyclic peptide as an inhibitor of Mycobacterium tuberculosis transcription: NMR and molecular dynamics simulations. Pharmaceuticals 2024, 17, 1545.

34. Prasad, A. K.; Samajdar, R.; Panwar, A. S.; Martin, L. L. Origin of secondary structure transitions and peptide self-assembly propensity in trifluoroethanol–water mixtures. The Journal of Physical Chemistry B 2024, 128, 7736–7749.

35. Xu, C.; Ren, Z.; Zhou, H.; Zhou, J.; Ho, C. P.; Wang, N.; Lee, C. Expanding chiral metamaterials for retrieving fingerprints via vibrational circular dichroism. Light: Science & Applications 2023, 12, 154.

36. Kuril, A. K.; Vashi, A.; Subbappa, P. K. A comprehensive guide for secondary and tertiary structure determination in peptides and proteins by circular dichroism spectrometer. Journal of Peptide Science 2025, 31, e3648.

37. Kumar, P.; Paterson, N. G.; Clayden, J.; Woolfson, D. N. De novo design of discrete, stable 310-helix peptide assemblies. Nature 2022, 607 (7918), 387–392.

38. Brown, R. A.; Marcelli, T.; De Poli, M.; Solà, J.; Clayden, J. Induction of unexpected left-handed helicity by an N-terminal L-amino acid in an otherwise achiral peptide chain. Angewandte Chemie 2012, 124 (6), 1424–1428.

39. Wei, Y.; Thyparambil, A. A.; Latour, R. A. Protein helical structure determination using CD spectroscopy for solutions with strong background absorbance from 190 to 230 nm. Biochimica et Biophysica Acta (BBA)-Proteins and Proteomics 2014, 1844 (12), 2331–2337.

40. Batra, A.; Darancet, P.; Chen, Q.; Meisner, J. S.; Widawsky, J. R.; Neaton, J. B.; Nuckolls, C.; Venkataraman, L. Tuning rectification in single-molecular diodes. Nano Letters 2013, 13 (12), 6233–6237.

