## Supplementary Information for "Role of stereochemistry on electron transport in peptides"

### S1. General Methods

#### Oligopeptide sequences

All oligopeptide sequences were purchased from GenScript (Piscataway, NJ). Mass spectrometry data for all the sequences are attached (**Supplementary Figures 1-10**). M, Y, and A denote the L-amino acids methionine, tyrosine, and alanine, respectively, whereas $Y_D$ and $A_D$ denote the corresponding D-amino acids.

#### Bulk spectroscopy

**UV-Vis spectroscopy.** UV–vis absorption spectra were recorded on an Agilent Cary 60 UV–Vis spectrophotometer at 298 K using a 1 cm quartz cuvette. The concentration of peptides was 0.1 mM.

**Fluorescence spectroscopy.** Fluorescence spectra were recorded on a Horiba FluoroMax-4 spectrofluorometer at 298 K using a 1 cm quartz cuvette. Both the excitation and emission slit widths were set to 5 nm. The peptide concentration was 0.1 mM.

**Circular dichroism (CD).** CD measurements were performed to characterize peptide secondary structure using a Jasco J1500 spectropolarimeter at 298 K. CD spectra were collected from 190 nm to 250 at 0.2 nm intervals, a rate of 50 nm/min, a response time of 2 s, a bandwidth of 1 nm, and 2 mm path length quartz cuvettes were used for solution samples. The peptide concentration was 0.1 mM.

**Two-dimensional nuclear magnetic resonance (2D NMR).** Diffusion-Ordered Spectroscopy (DOSY) NMR was measured on Agilent VNS 750. Nuclear Overhauser Effect Spectroscopy (NOESY) NMR was measured on Carver B500 Bruker Advance III HD.

#### Single-molecule conductance measurements

Single-molecule conductance measurements were performed using a custom-built scanning tunneling microscope break junction (STM-BJ).[1–6] Gold substrates were prepared by evaporating 100 nm of gold onto polished Ted Pella AFM specimen disks with an e-beam evaporator. STM tips were prepared with 0.25 mm Au wire (99.998%, Alfa Aesar). All STM-BJ experiments were carried out in Corning™ cell culture grade water. Due to the polarity of the solvent, STM tips were coated with an Apiezon wax to prevent Faradaic currents from masking characteristic molecular features.[7] During experiments, the STM tip is controlled by a piezoelectric micro-positioner to repeatedly form and break molecular junctions, and the current was recorded and analyzed during this process. A variable-gain low noise current amplifier (DLPCA-200 from Artisan Technology Group) was used to accurately convert current to voltage for data processing. Conductance histograms (determined from > 5000 individual molecules per experiment) are generated for all molecules without data selection. The 1D conductance histograms were generated on a logarithmic scale [$\log(G/G_0)$] from −5.5 to 0 with a bin size of 0.01 and subsequently smoothed using a 1D Gaussian filter with a standard deviation of 20 bins. The 2D molecular conductance histograms show the distribution of conductance values together with junction separation distances, providing insights into the evolution of conductance as the junction is extended.

**Molecular dynamics (MD) simulations**

MD simulations were performed to generate conformational ensembles for the MAAAM, MYYYM, MYY$_D$YM, and MAA$_D$AM molecular junctions at three anchor displacements (referred to as stages 6 Å, 9 Å, and 12 Å). For each peptide, 16 initial structures were prepared using the PeptideBuilder python package.[8] Phi and psi backbone dihedral angles of each of the 16 structures were randomized independently. Each backbone dihedral angle was initialized to a random value between -180 and 180 degrees. Hydrogens were added to the peptides with the VMD plugin PSFGEN[9] and we used the NTER and CTER terminal patches to create positively and negatively charged N- and C-termini, respectively. Peptide structures were then solvated in a cubic box of TIP3P water of side length 38 Å using the VMD SOLVATE plugin.[9] The solvated systems were then subjected to MD simulations with the CHARMM36m protein force field[10,11] using OpenMM 7.7.0.[12] Dynamics were integrated using the LangevinMiddleIntegrator[13] with friction coefficient of 1 ps$^{-1}$, temperature of 300 K, and a timestep of 4 fs. Hydrogen mass repartitioning was not utilized. Bonds involving hydrogen atoms, and all bonds and angles involving water were constrained.[13] Nonbonded interactions were computed using a cutoff of 12 Å with smooth switching starting at 10 Å. Electrostatic interactions were evaluated using particle mesh Ewald[14] (PME) summation with error tolerance of 0.0005. Each of the 16 replicates for each of the three holding stages was simulated for 200 ns, for a total aggregate simulation time of 38.4 μs (4 peptides × 3 stages × 16 replicates × 200 ns). The last 190 ns of each simulation was used for subsequent analysis.

Holding stages were enforced using a series of custom external potentials, applied using OpenMM's custom force classes, as described in our prior work.[4] A series of custom potentials were implemented to implicitly represent interactions between the peptide and gold particles. Three potentials were defined: (1) a potential to restrain the distance between the anchors of the molecular junction along the pulling axis to 6 Å, 9 Å, or 12 Å (representing the restraints imposed by connections to the gold electrodes); (2) a per-atom charge-dependent potential along the pulling axis accounting representing electric field forces arising from a voltage-biased junction; and, (3) a potential that orients methionine's thioether moiety such that the average position of each sulfur's lone pairs is oriented towards the (implicitly represented) gold electrodes along the pulling axis. These potentials are described in detail in the next section and depicted in **Supplementary Figure 24**.

A key challenge for simulating single-molecule pulling processes is the large difference between the pulling rates used in experiments and those accessible by MD simulations. Typical experimental pulling rates are on the order of Ångstroms per millisecond (1 Å per 5 ms in the present study), whereas single-trajectory MD simulations (at most) typically reach ms timescales, e.g., with the use of bespoke hardware[15] or massively distributed computing schemes.[16] In addition, the need for multiple independent simulation replicas to claim ensemble convergence and statistical certainty of key observables further restricts simulations to sub-experimental timescales. However, because the experimental pulling rate is also slow relative to characteristic relaxation timescales of small peptides, we assume that all molecular conformations accessible at a given end-to-end distance are sampled during each step of the experimental pulling process. In other words, experimental pulling occurs as an equilibrium process. Rather than performing costly simulations of the entire pulling process, it is computationally more feasible to simulate the molecular junction at various holding (end-to-end distance) stages representing the different separation distances arising during the pulling experiments.

Using this approach, we performed a series of independent simulations where we restrained the end-to-end (sulfur-sulfur) distance along the pulling axis to one of three distances spanning the range of end-to-end distances (6 Å, 9 Å, or 12 Å). We define the pulling axis as the *z*-axis in our simulations. Schematic illustrations for each potential are shown in **Supplementary Figure 24**. The functional form of the potential utilized to enforce this restraint is given in Equation 1:

$$U_1 = \frac{1}{2} k_1 [(z_{S_2} - z_{S_1}) - z_0]^2, \tag{1}$$

where the coefficient $k_1$ is the force constant of the harmonic potential, $z_{S1}$ and $z_{S2}$ are the *z*-coordinates of the sulfur atoms of the N- and C-terminal methionine residues respectively, and $z_0$ is the equilibrium distance for a given stage. We use a value of 1 kcal/mol/Å$^2$ for $k_1$ and utilize three independent holding stages with $z_0$ equal to either 6 Å, 9 Å, or 12 Å. This force constant was selected such that the resulting distributions of $z_{S2} - z_{S1}$ distances have slight overlap (**Supplementary Figures 24a,d**).

By restraining the *z*-displacement between the sulfur atoms, rather than the distance, the movement of each sulfur atom is effectively restrained to one of two parallel planes which implicitly represent two parallel planes of gold electrode.

A potential is introduced to represent an applied electric field due to the voltage difference across the two electrodes. The functional form is given in Equation 2:

$$U_2 = \sum_{i=1}^{N_{atoms}} -q_i E z_i = \sum_{i=1}^{N_{atoms}} -q_i \left(\frac{V}{z_0 + 2l_{S-Au}}\right) z_i \tag{2}$$

where $N_{atoms}$ is the total number of atoms in each system including solvent, $q_i$ is the charge of atom *i*, $z_i$ is the z-coordinate of atom *i*, $z_0$ is the equilibrium end-to-end distance (displacement along *z*) for a holding stage, and $l_{S-Au}$ is the length of the sulfur-gold bond.

We further introduce a potential to orient each sulfur atom's lone pairs in either the positive or negative *z*-direction, such that a feasible dative bond may occur between the sulfur and a fictitious gold particle. This is a key step in ensuring that any conformation generated by MD simulations can be placed into a gold-gold junction for subsequent NEGF-DFT calculations. Because electron lone pairs are not explicitly represented in atomistic MD simulations, we define surrogate vectors that involve each sulfur's adjacently bonded carbon atoms to act as a proxy for the direction of the electron lone pairs (**Supplementary Figure 24b**). We impose a restraint directly on the dot product of each surrogate vector with the pulling axis. The functional form of this potential is shown in Equation 3:

$$U_3 = \sum_{i=1}^{2} k_3 \left[ \left| \vec{r}_{S_i} - \left( \frac{\vec{r}_{CG_i} - \vec{r}_{CE_i}}{2} \right) \right| \cdot \vec{z} \right] = \sum_{i=1}^{2} k_3 [|\vec{p}_i| \cdot \vec{z}](-1)^i \quad (3)$$

where $\vec{r}_{S_i}$ represents the three-dimensional Cartesian coordinates of the sulfur atom of interest, with $S_1$ and $S_2$ subscripts indicating the sulfur atoms in the N- and C-terminal methionine residues, respectively, $\vec{r}_{CG_i}$ and $\vec{r}_{CE_i}$ are Cartesian coordinates of the adjacent carbon atoms covalently bonded to each sulfur of interest, and $\vec{z}$ is the unit vector in the direction of the *z*-axis. Vertical lines denote vector normalization. The final term in the equation determines the sign of the potential (and thus the direction of the surrogate vector) allowing for one sulfur's lone pair to be oriented in the positive *z*-direction while the other in the negative *z*-direction. The value of $k_3$ is taken as 10 kcal/mol, resulting in a strong potential that tightly secures the orientation of sulfur lone pairs towards the implicitly represented gold electrodes (**Supplementary Figures 24c,e**).

**Density functional theory (DFT) calculations**

Single-point density functional theory (DFT) calculations were performed using ORCA (version 6.0.0) on a representative molecular conformation extracted from molecular dynamics (MD) simulations. Electronic energies were evaluated at the ωB97X-3c level of theory. Self-consistent field (SCF) convergence was enforced using TightSCF criteria. Molecular orbital (MO) information was obtained from the converged wavefunction, and orbital populations were printed using the Print[P_MOs] keyword. Orbital visualizations were performed for the molecular junction model comprising the molecule of interest with one gold atom on each side. Frontier molecular orbitals (HOMO-1, HOMO, LUMO, and LUMO+1) were visualized using an isosurface value of 0.02 a.u. The resulting orbital energies and populations were used for subsequent projected density of states (PDOS) analysis.

**Nonequilibrium Green's function-density functional theory (NEGF-DFT)**

Representative molecular conformations generated of the most probable conformations by MD were used in quantum mechanics (QM) calculations to enable direct comparison between theory and experimental results. Non-equilibrium Green's function-density functional theory (NEGF-DFT) calculations for MYYYM, $MYY_DYM$, MAAAM, and $MAA_DAM$ were performed with double-zeta (DZ) basis sets for gold atoms and double-zeta polarized (DZP) basis sets for carbon, hydrogen, oxygen, sulfur, and nitrogen. NEGF-DFT calculations were performed with a DFT-based non-equilibrium Green's function (NEGF) approach using the TranSiesta and Tbtrans package.[17–19] Electrode configurations contain 8 layers of 16 gold atoms along with a pyramid of 9 Au atoms. Sulfur atoms in the peptoids were made to interact with the gold atoms using a trimer binding motif, as previously described[4]. Geometry relaxation of the sequences were performed using generalized gradient approximation-Perdew-Burke-Ernzerhof (GGA-PBE) functional[20] using the TranSiesta package.[18]

Electrode calculations were carried out with a $4 \times 4 \times 50$ k-mesh. The geometry relaxation was carried out using a $4 \times 4 \times 1$ k-mesh, which was performed till all the forces were < 0.05 eV/Å. After the junction was relaxed, the transport calculations were carried out using the TranSiesta package[17,19] with the same functionals, basis sets, pseudopotential, and k-mesh as the geometry relaxation. Convergence was tested prior to transmission calculations, using a real axis integration internal from -40 eV to infinity[17]; this includes a crossing in the imaginary axis at 2.5 eV, and the $\gamma$ value is $-10k_{\mathrm{B}}T$. The circle grid consists of 102 Gauss-Legendre points, and 15 Gauss-Fermi points for the tail portion. Tbtrans[19] was used to carry out the NEGF calculations and to obtain electron transmission as a function of energy (relative to the fermi energy level). The transmission plots are shifted with respect to the Fermi energy values of each peptide. NEGF calculations were carried out from -3 eV to 3 eV with 0.05 eV energy increments.

### S2. Mass spectrometry data

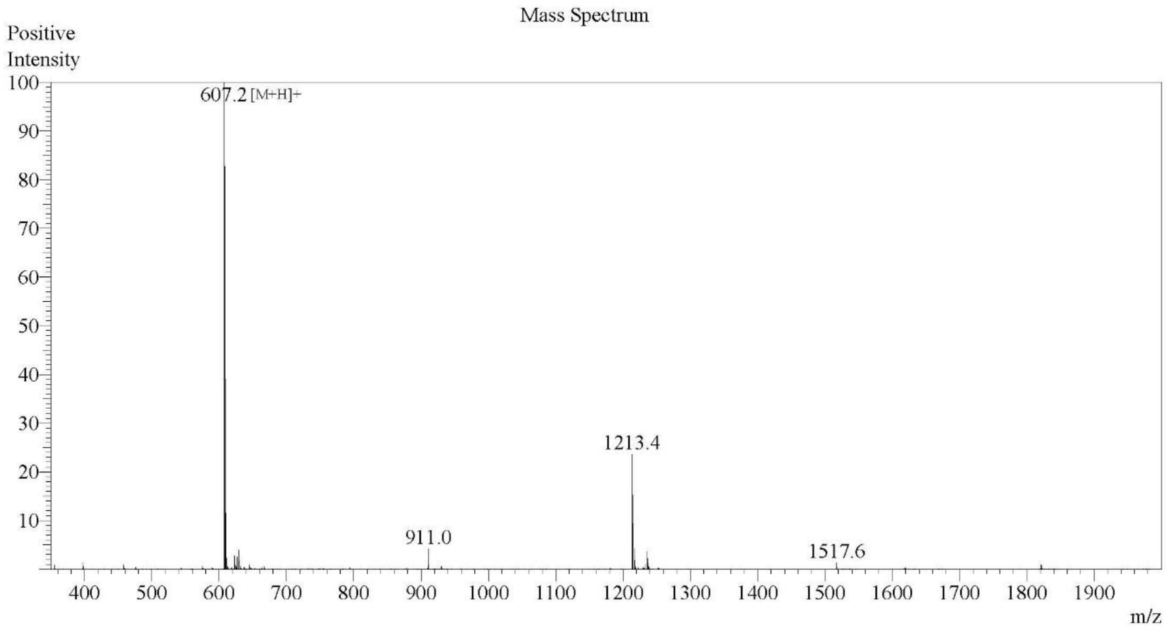

**Supplementary Figure 1:** Electrospray ionization (ESI) mass spectrometry data for peptide sample MYYM. Theoretical molecular weight is 606.76 m/z. Observed molecular weight is 606.2 m/z.

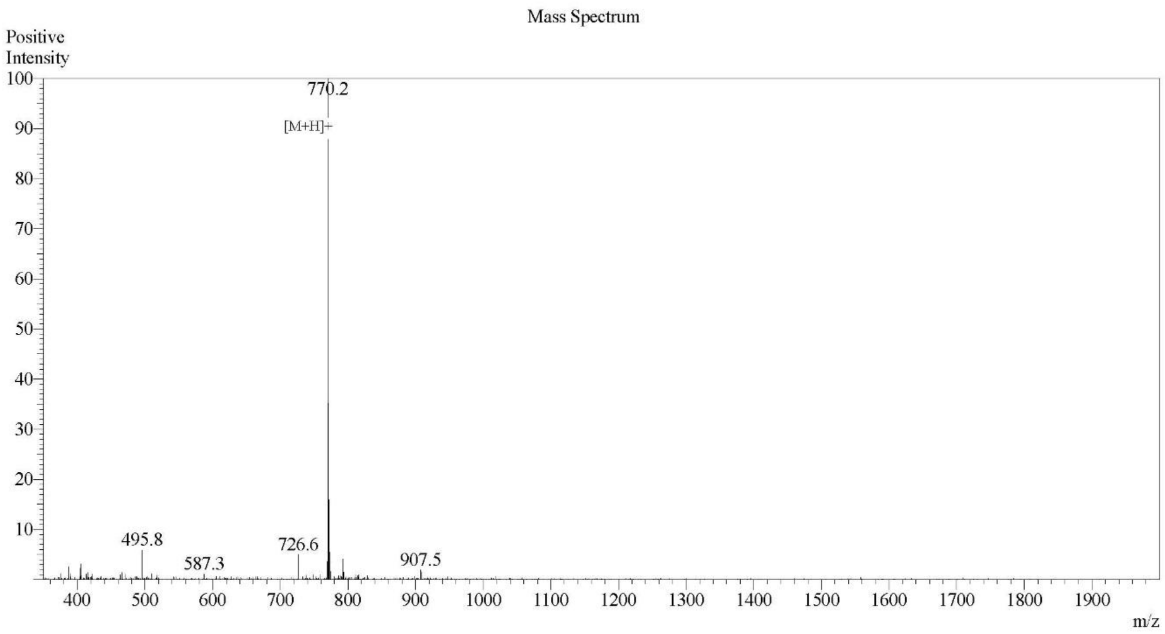

**Supplementary Figure 2:** Electrospray ionization (ESI) mass spectrometry data for peptide sample MYYYM. Theoretical molecular weight is 769.93 m/z. Observed molecular weight is 769.2 m/z.

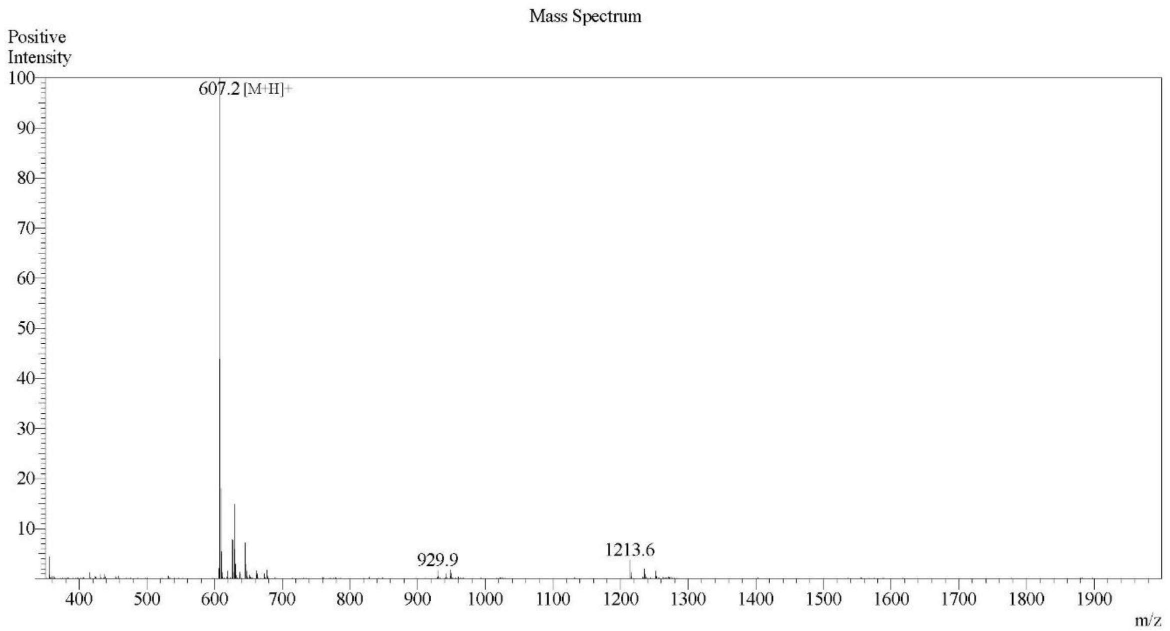

**Supplementary Figure 3:** Electrospray ionization (ESI) mass spectrometry data for peptide sample MYY$_D$M. Theoretical molecular weight is 606.76 m/z. Observed molecular weight is 606.2 m/z.

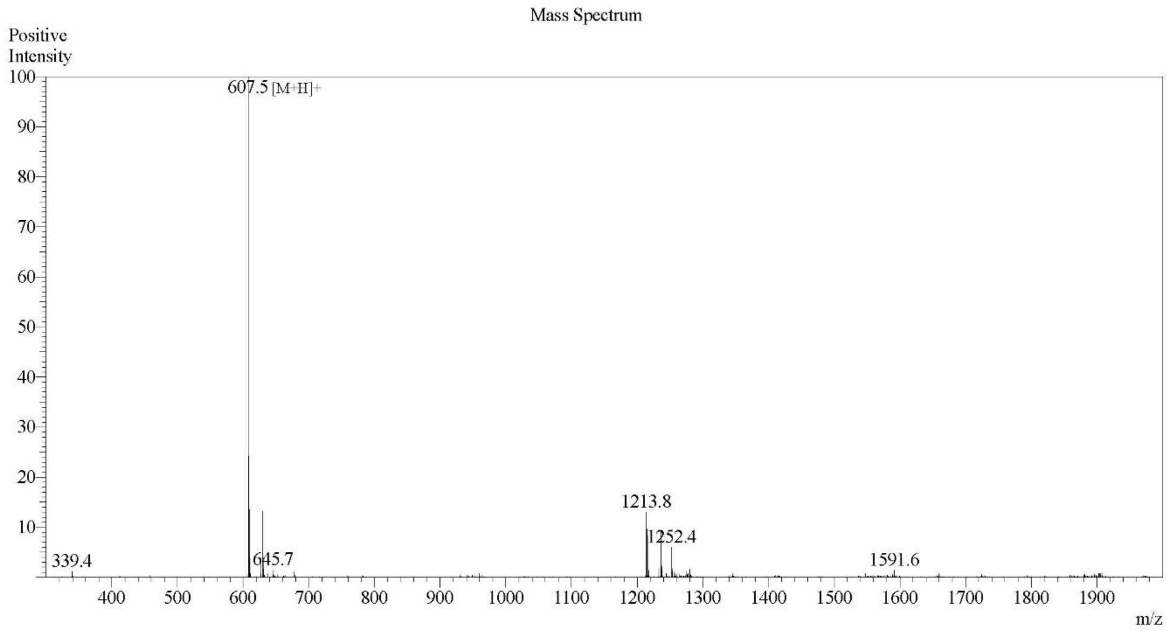

**Supplementary Figure 4:** Electrospray ionization (ESI) mass spectrometry data for peptide sample $MY_DYM$. Theoretical molecular weight is 606.76 m/z. Observed molecular weight is 606.5 m/z.

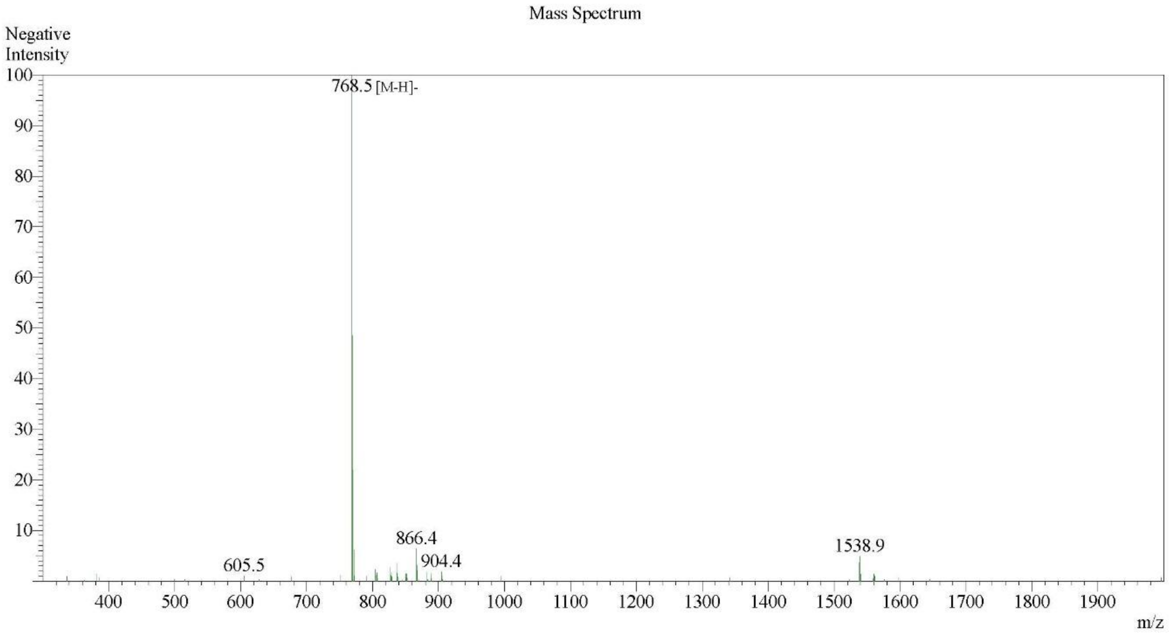

**Supplementary Figure 5:** Electrospray ionization (ESI) mass spectrometry data for peptide sample MYY$_D$YM. Theoretical molecular weight is 769.93 m/z. Observed molecular weight is 769.5 m/z.

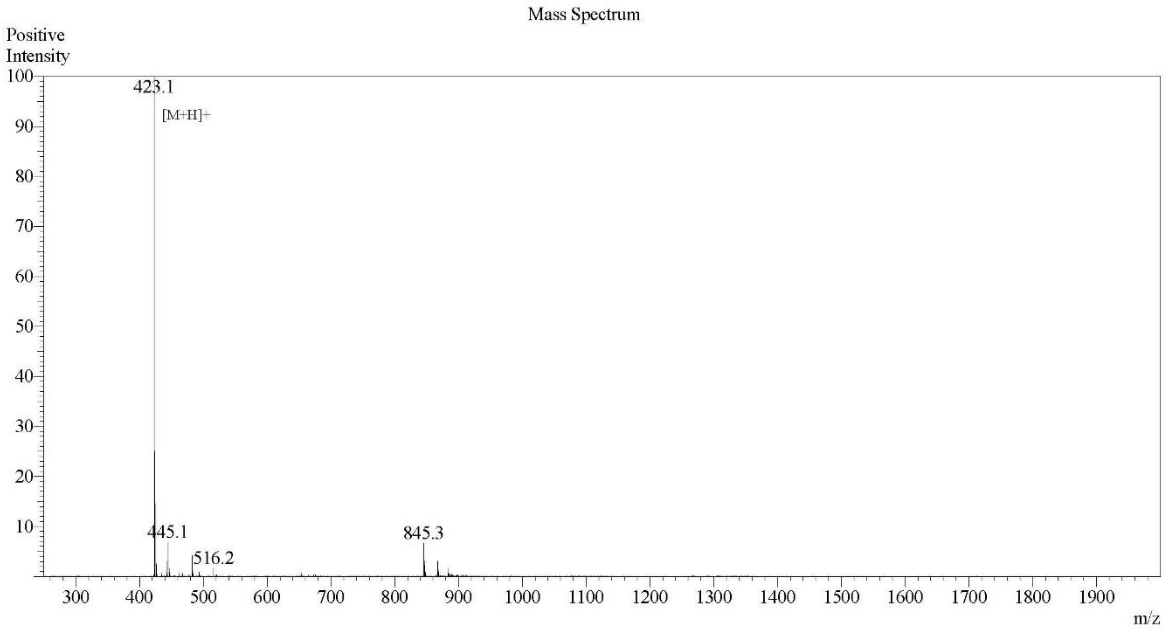

**Supplementary Figure 6:** Electrospray ionization (ESI) mass spectrometry data for peptide sample MAAM. Theoretical molecular weight is 422.57 m/z. Observed molecular weight is 422.1 m/z.

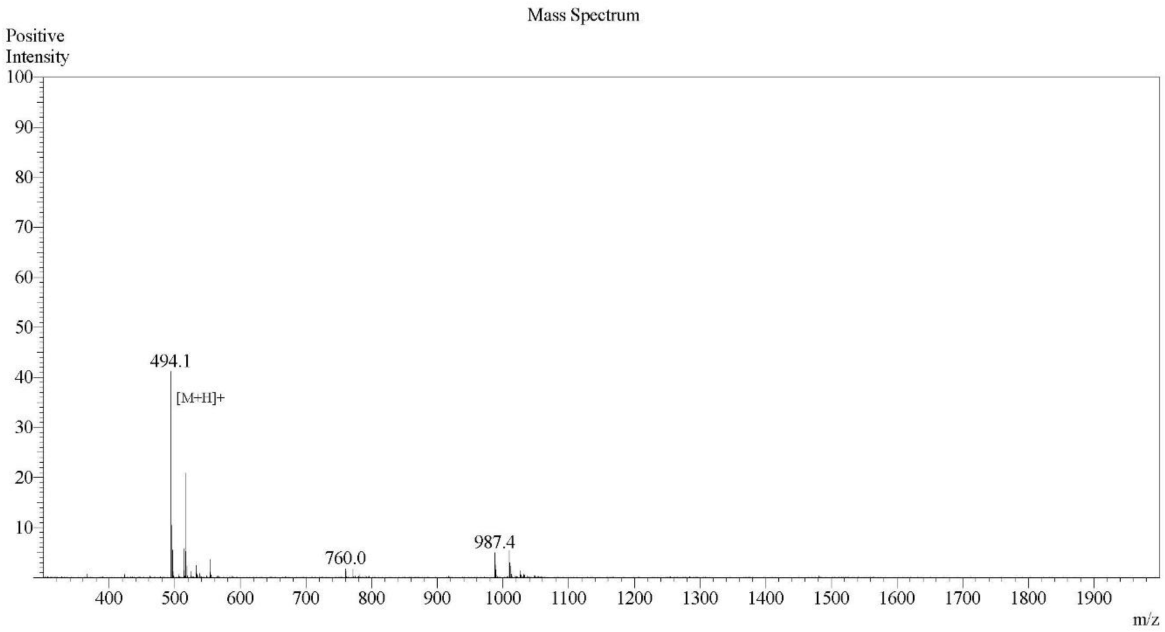

**Supplementary Figure 7:** Electrospray ionization (ESI) mass spectrometry data for peptide sample MAAAM. Theoretical molecular weight is 493.64 m/z. Observed molecular weight is 493.1 m/z.

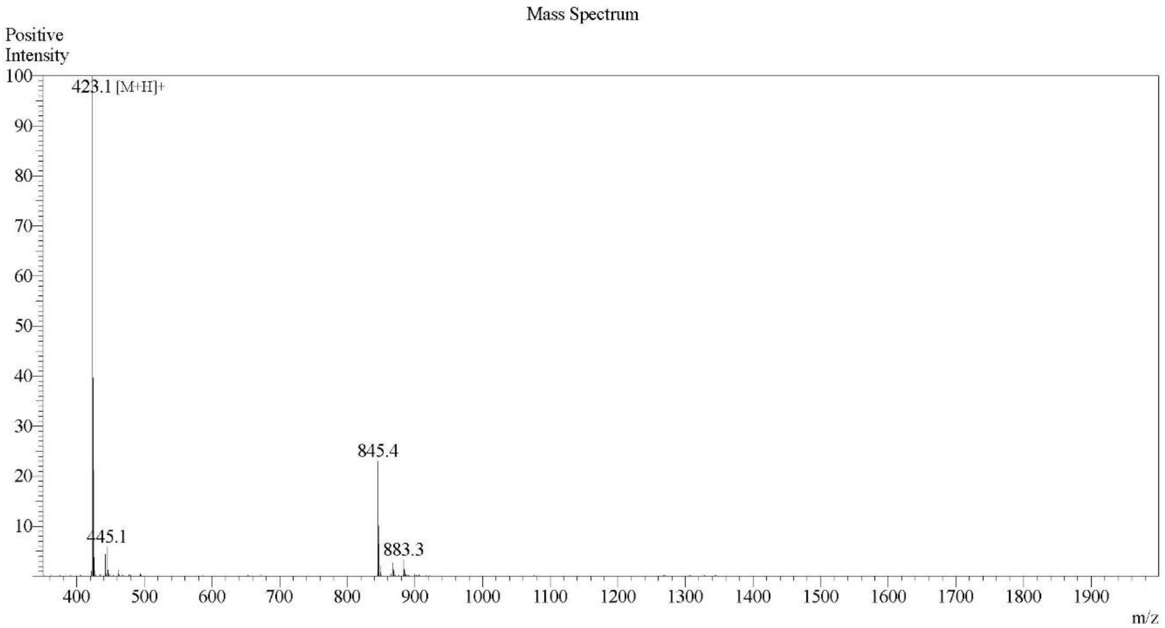

**Supplementary Figure 8:** Electrospray ionization (ESI) mass spectrometry data for peptide sample MA$_D$AM. Theoretical molecular weight is 422.57 m/z. Observed molecular weight is 422.1 m/z.

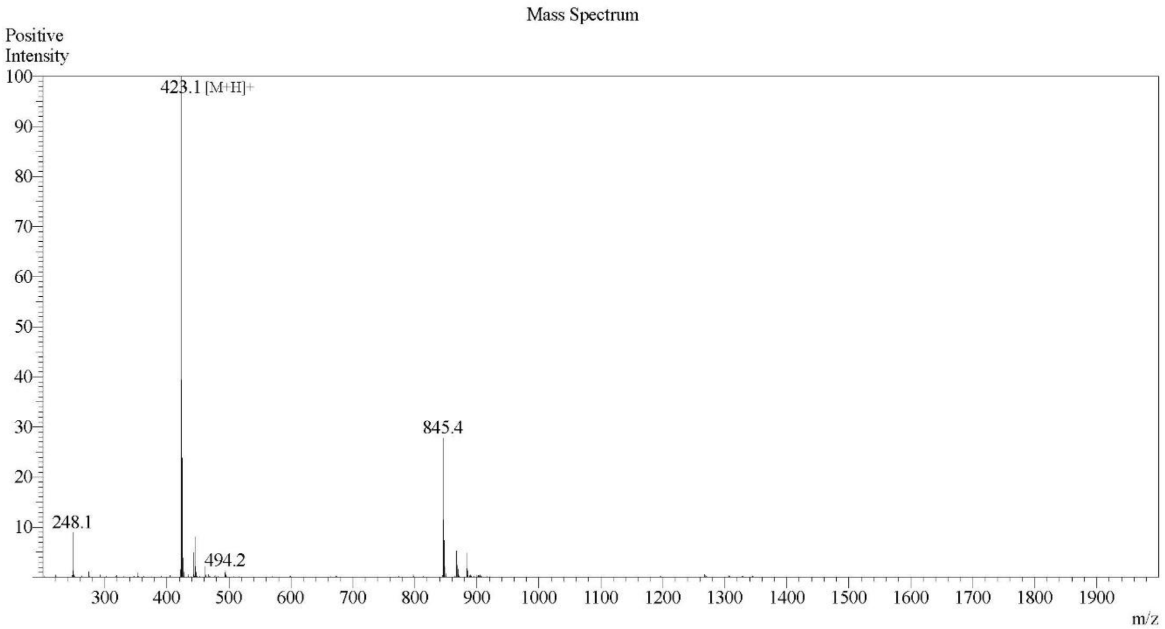

**Supplementary Figure 9:** Electrospray ionization (ESI) mass spectrometry data for peptide sample MAA$_D$M. Theoretical molecular weight is 422.57 m/z. Observed molecular weight is 422.1 m/z.

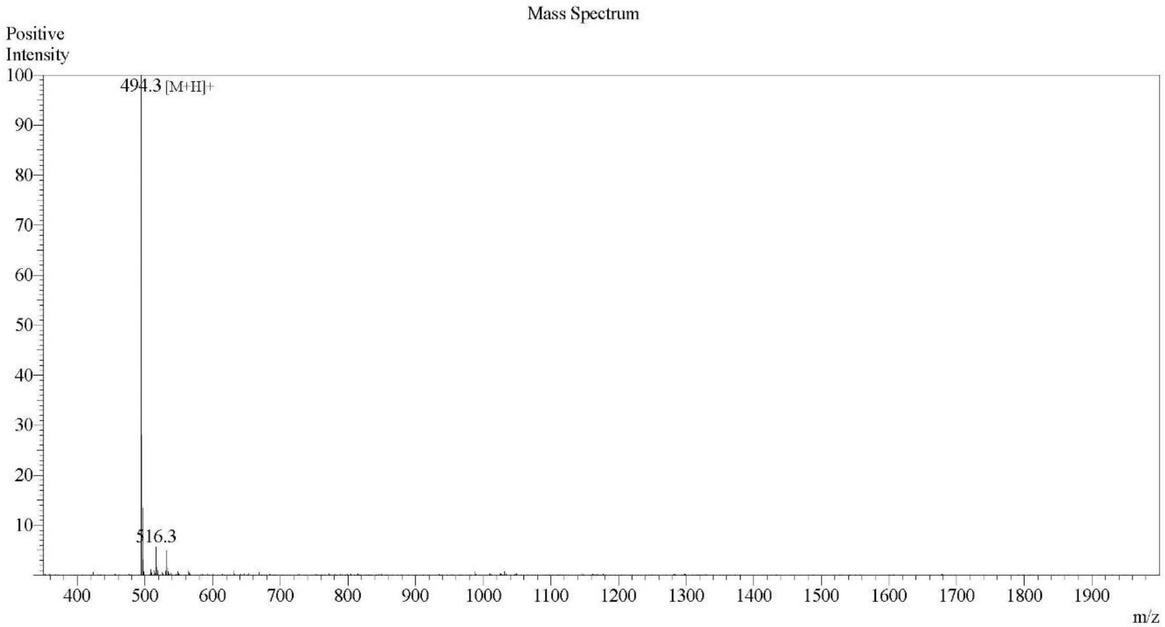

**Supplementary Figure 10:** Electrospray ionization (ESI) mass spectrometry data for peptide sample MAA$_D$AM. Theoretical molecular weight is 493.64 m/z. Observed molecular weight is 493.1 m/z.

**S3. UV-visible and fluorescence spectroscopy**

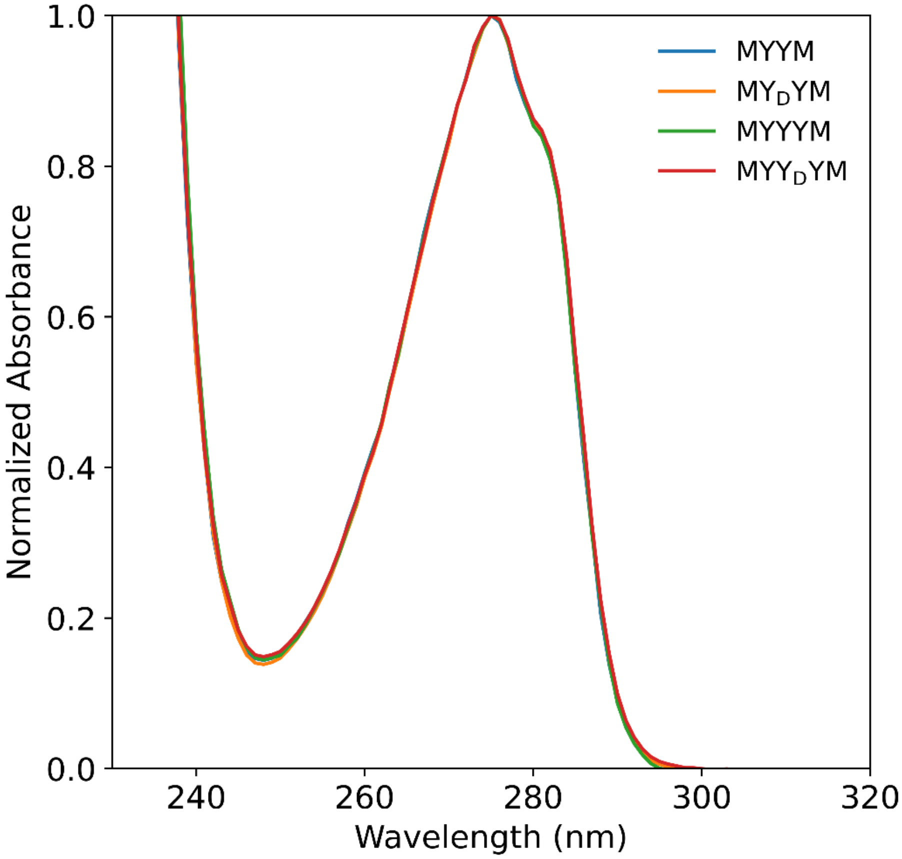

**Supplementary Figure 11:** UV-visible spectroscopy data for tyrosine-based peptide sequences in water. A distinct peak is observed at 275 nm, accompanied with shoulders at 282 nm. The concentration of the peptides was 0.1 mM.

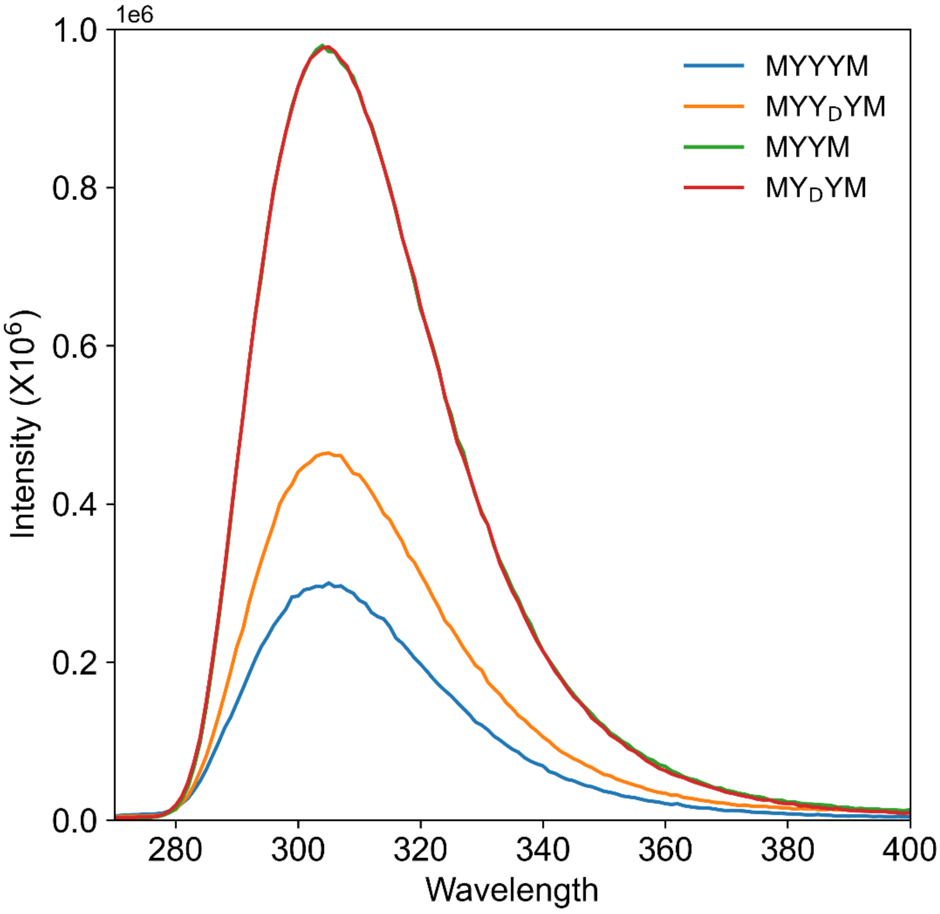

**Supplementary Figure 12:** Fluorescence spectroscopy for tyrosine-based peptide sequences in water. All peptide sequences show an emission peak at 305 nm. Spectra for peptide sequences $MY_DYM$ (red) and MYYM (green) are superimposed. The concentration of the peptides was 0.1 mM.

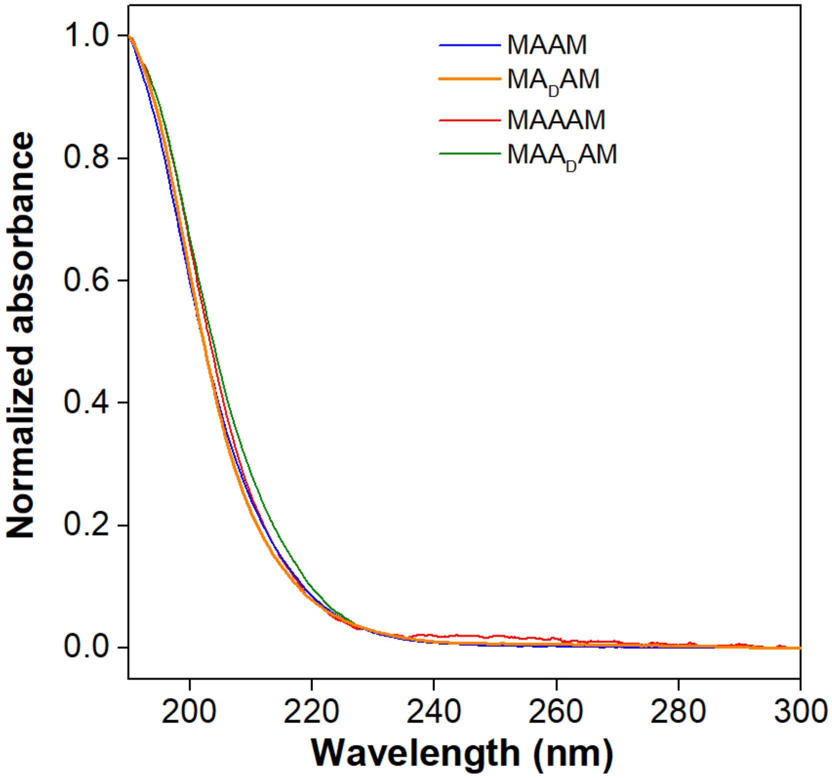

**Supplementary Figure 13:** UV-visible spectroscopy data for alanine-based sequences in water. No distinct peaks are observed in the aromatic region. Absorption near 190–220 nm is expected due to C=O transitions of the peptide backbone.[21,22] The concentration of the peptides was 0.1 mM.

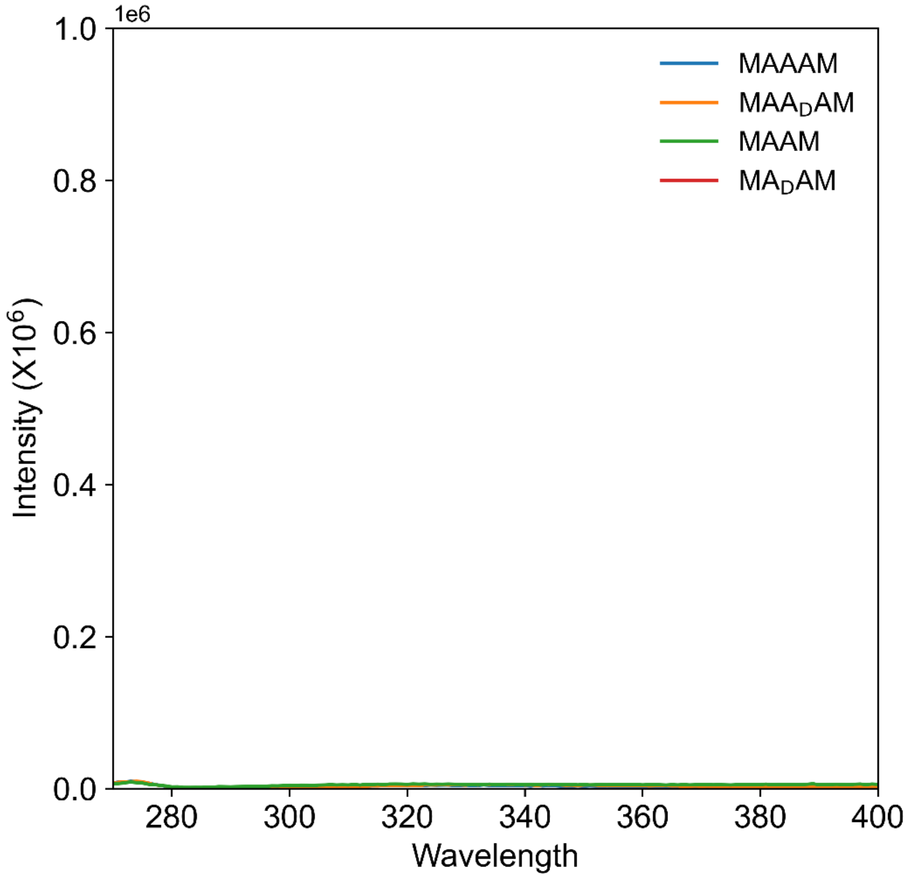

**Supplementary Figure 14:** Fluorescence spectroscopy for alanine-based sequences in water. No significant peaks are observed for any of the alanine-based sequences. The concentration of the peptides was 0.1 mM.

**S4. Two-dimensional (2D) nuclear magnetic resonance (NMR) spectroscopy**

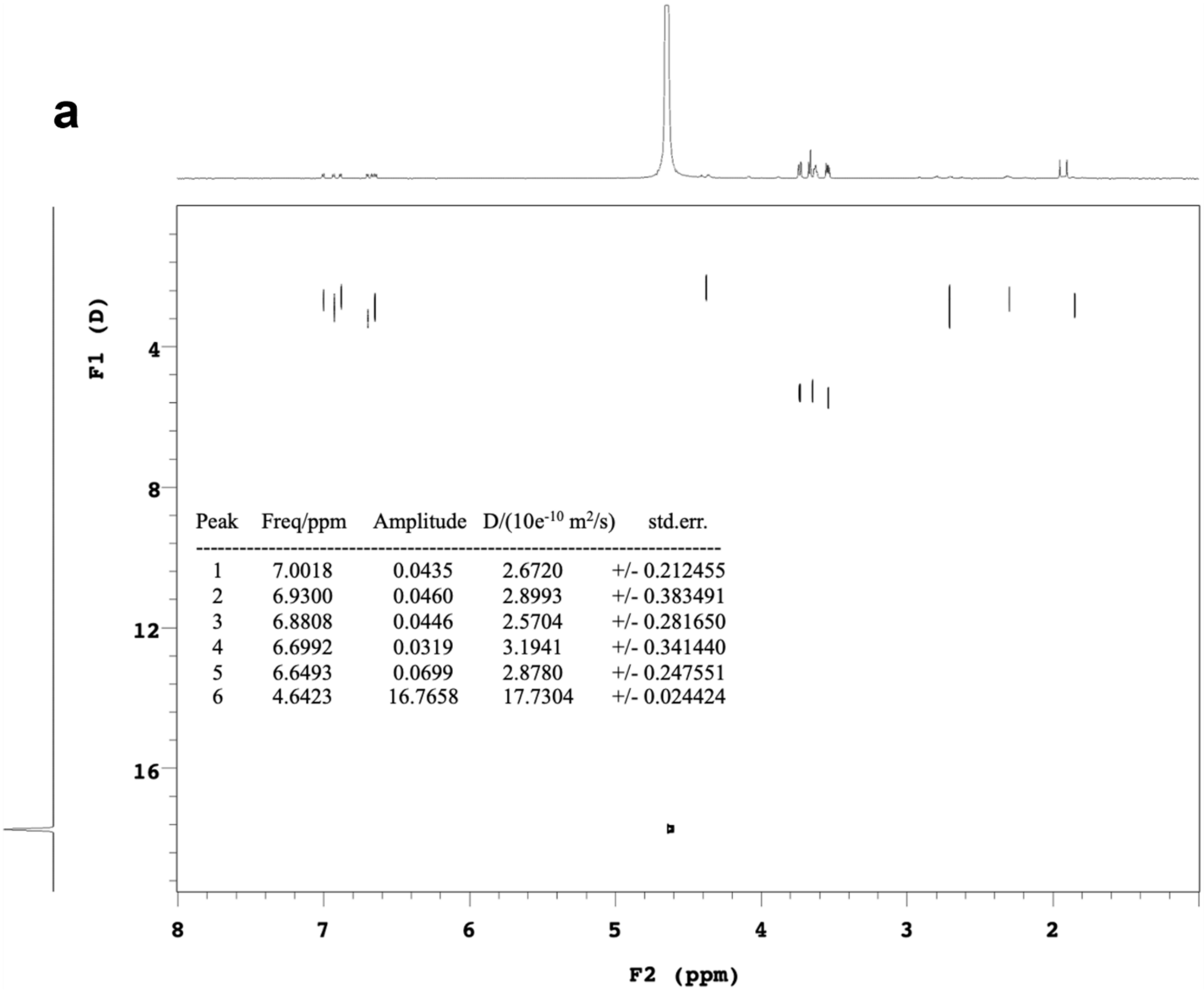

| Peak | Freq/ppm | Amplitude | D/(10e$^{-10}$ m$^2$/s) | std.err. |
|---|---|---|---|---|
| 1 | 7.0018 | 0.0435 | 2.6720 | +/- 0.212455 |
| 2 | 6.9300 | 0.0460 | 2.8993 | +/- 0.383491 |
| 3 | 6.8808 | 0.0446 | 2.5704 | +/- 0.281650 |
| 4 | 6.6992 | 0.0319 | 3.1941 | +/- 0.341440 |
| 5 | 6.6493 | 0.0699 | 2.8780 | +/- 0.247551 |
| 6 | 4.6423 | 16.7658 | 17.7304 | +/- 0.024424 |

**Supplementary Figure 15a.** Diffusion-ordered spectroscopy (DOSY) NMR for MYYYM Deuterium oxide ($D_2O$) was used as the solvent. The concentration of the peptides was 0.1 mM.

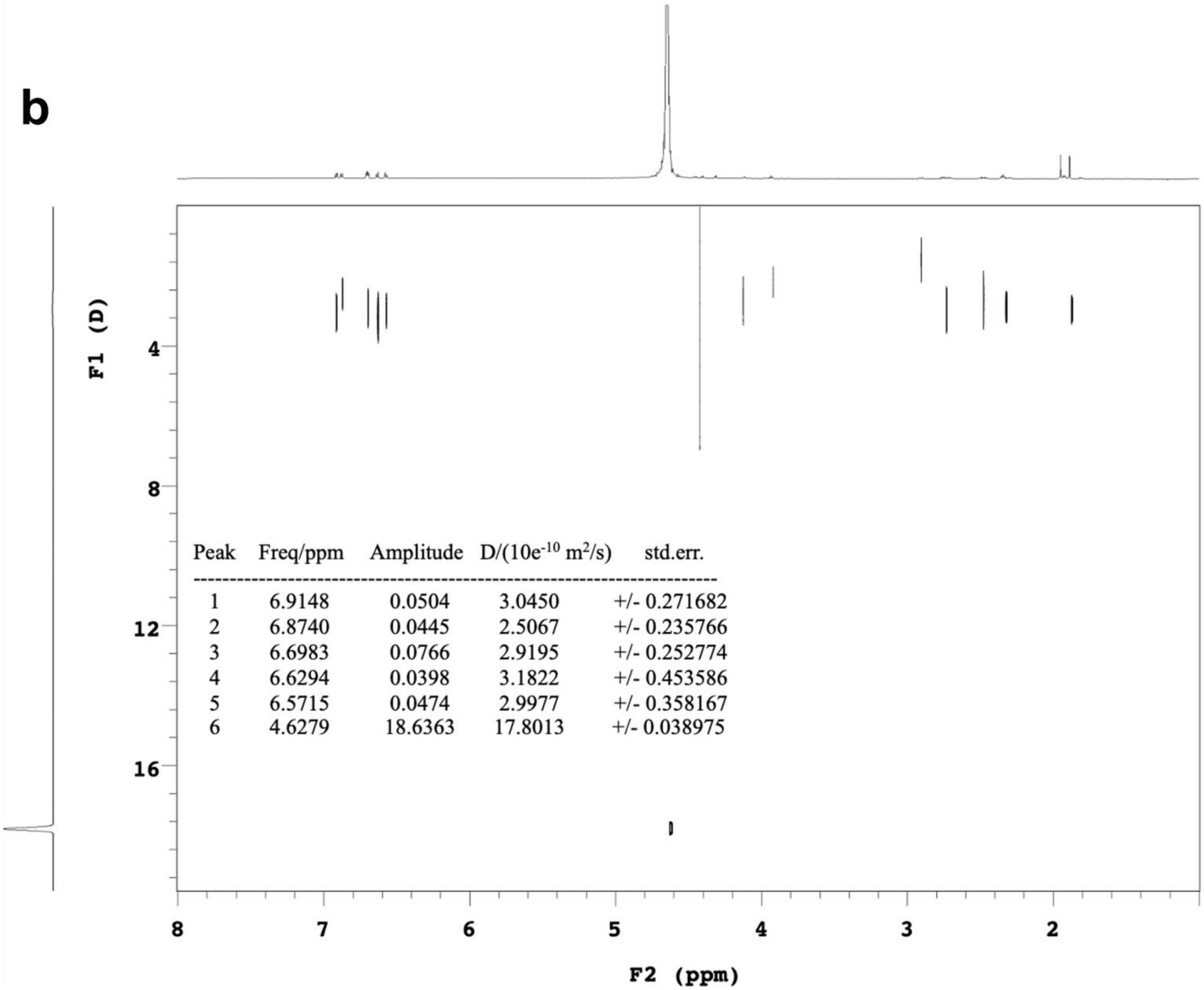

**Supplementary Figure 15b.** Diffusion-ordered spectroscopy (DOSY) NMR for MYY$_D$YM. Deuterium oxide ($D_2O$) was used as the solvent. The concentration of the peptides was 0.1 mM.

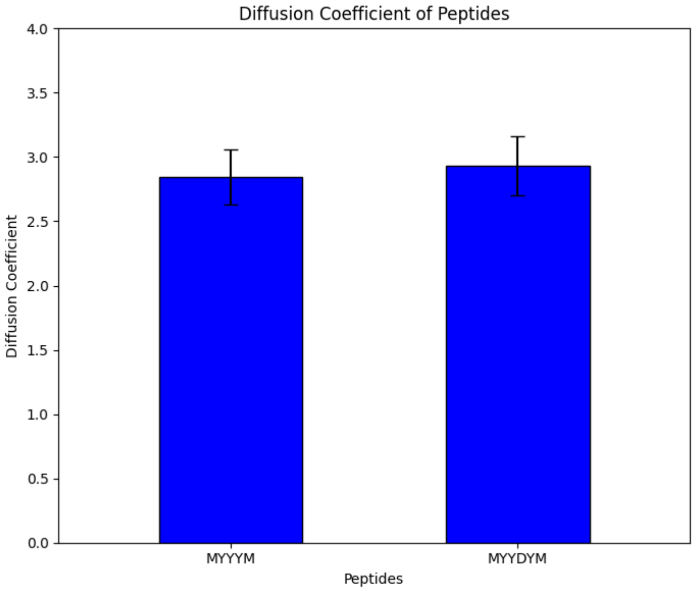

**Supplementary Figure 16:** Diffusion-ordered spectroscopy (DOSY) NMR for peptides MYYYM and MYY$_D$YM indicating similar diffusion coefficients. These results indicate similar molecular size and shape for the tyrosine-based sequences in solution phase. Only peaks from phenyl rings (1 to 5) were used to calculate average diffusion coefficients of each molecule. The diffusion coefficients (x $10^{-10}$ $m^2/s$) of MYYYM and MYY$_D$YM are 2.8428 and 2.9032, respectively. The concentration of the peptides was 0.1 mM.

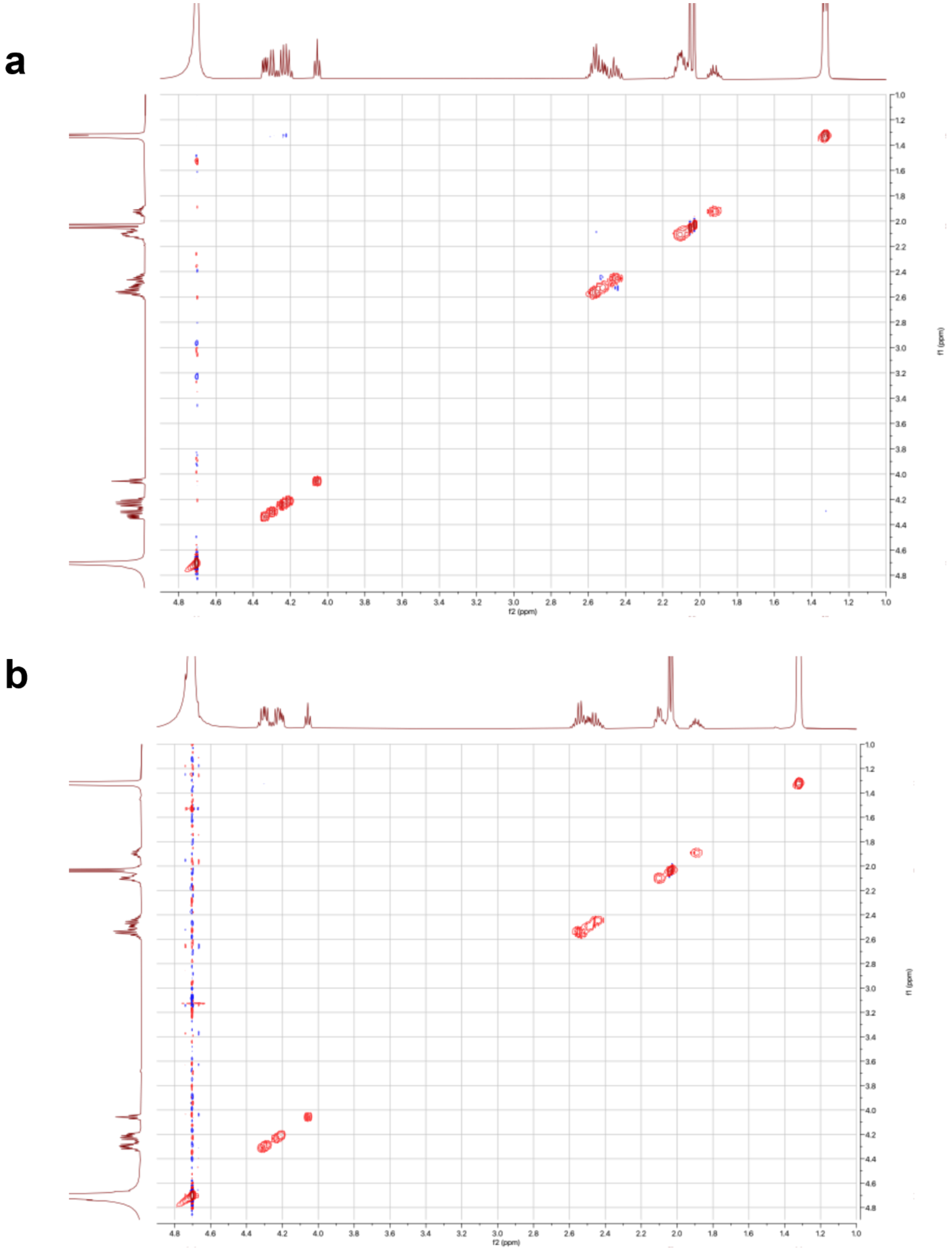

**Supplementary Figure 17:** Nuclear Overhauser effect spectroscopy (NOESY) NMR for peptides (a) MAAAM and (b) $MAA_DAM$. Deuterium oxide ($D_2O$) was used as the solvent. The concentration of the peptides was 0.1 mM.

### S5. Circular Dichroism (CD)

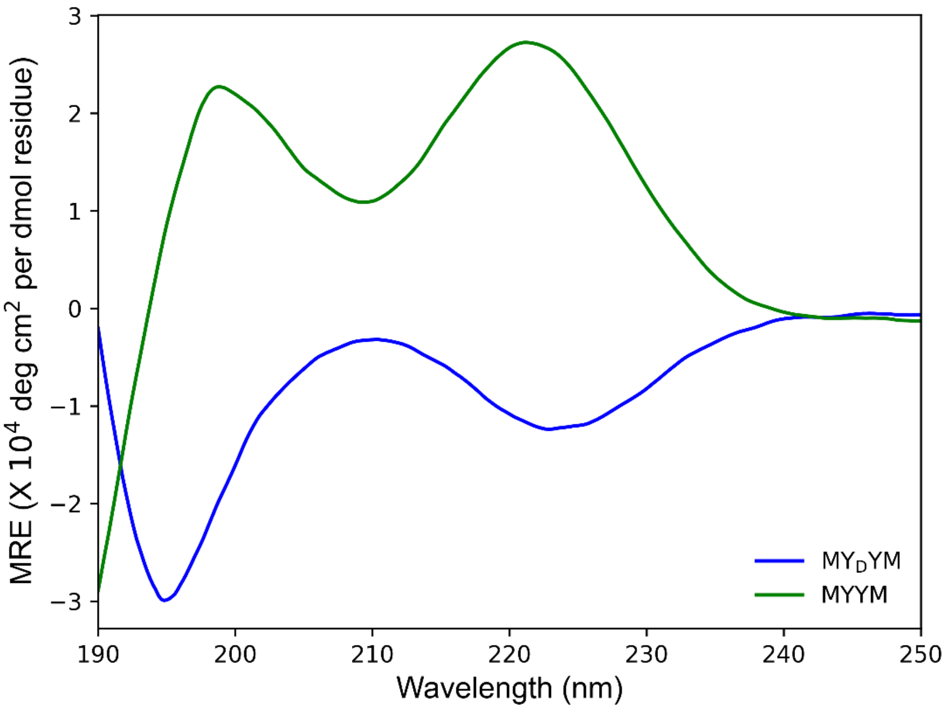

**Supplementary Figure 18:** Circular dichroism data for tyrosine-based tetrapeptides in water. The concentration of the peptides was 0.1 mM.

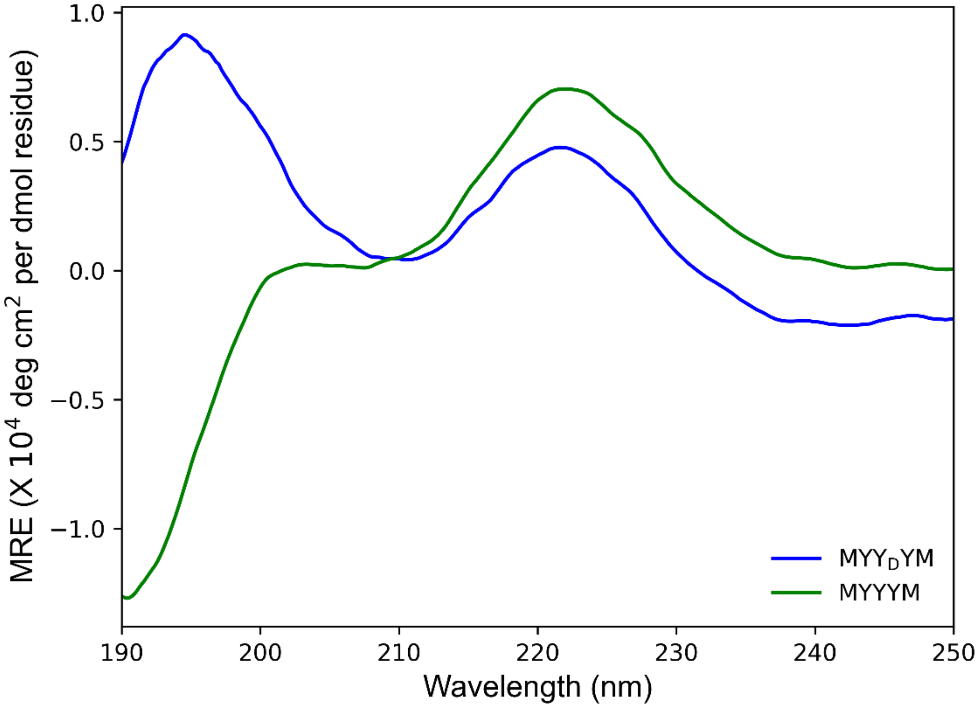

**Supplementary Figure 19:** Circular dichroism data for tyrosine-based pentapeptides in water. The concentration of the peptides was 0.1 mM.

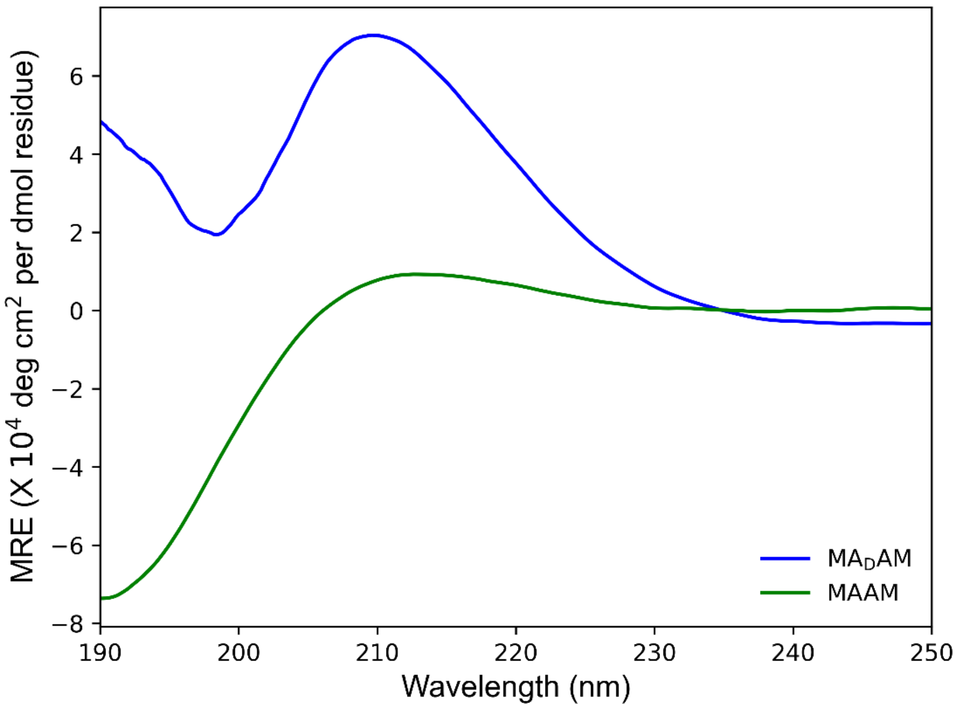

**Supplementary Figure 20:** Circular dichroism data for alanine-based tetrapeptides in water. The concentration of the peptides was 0.1 mM.

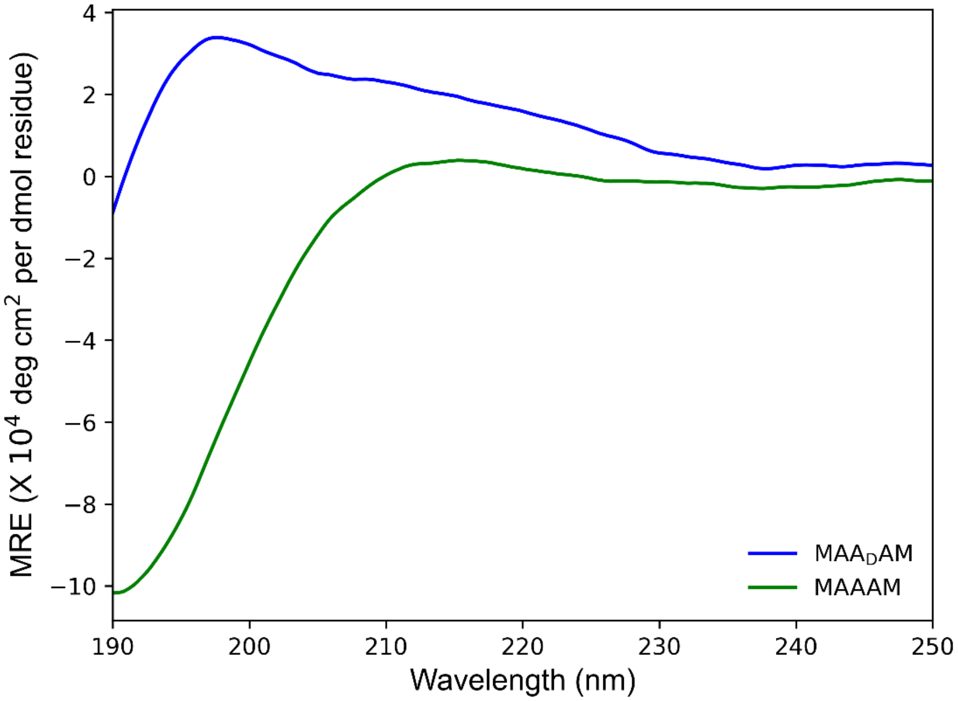

**Supplementary Figure 21:** Circular dichroism data for alanine-based pentapeptides in water. The concentration of the peptides was 0.1 mM.

### S6. Single-molecule electronic experiments

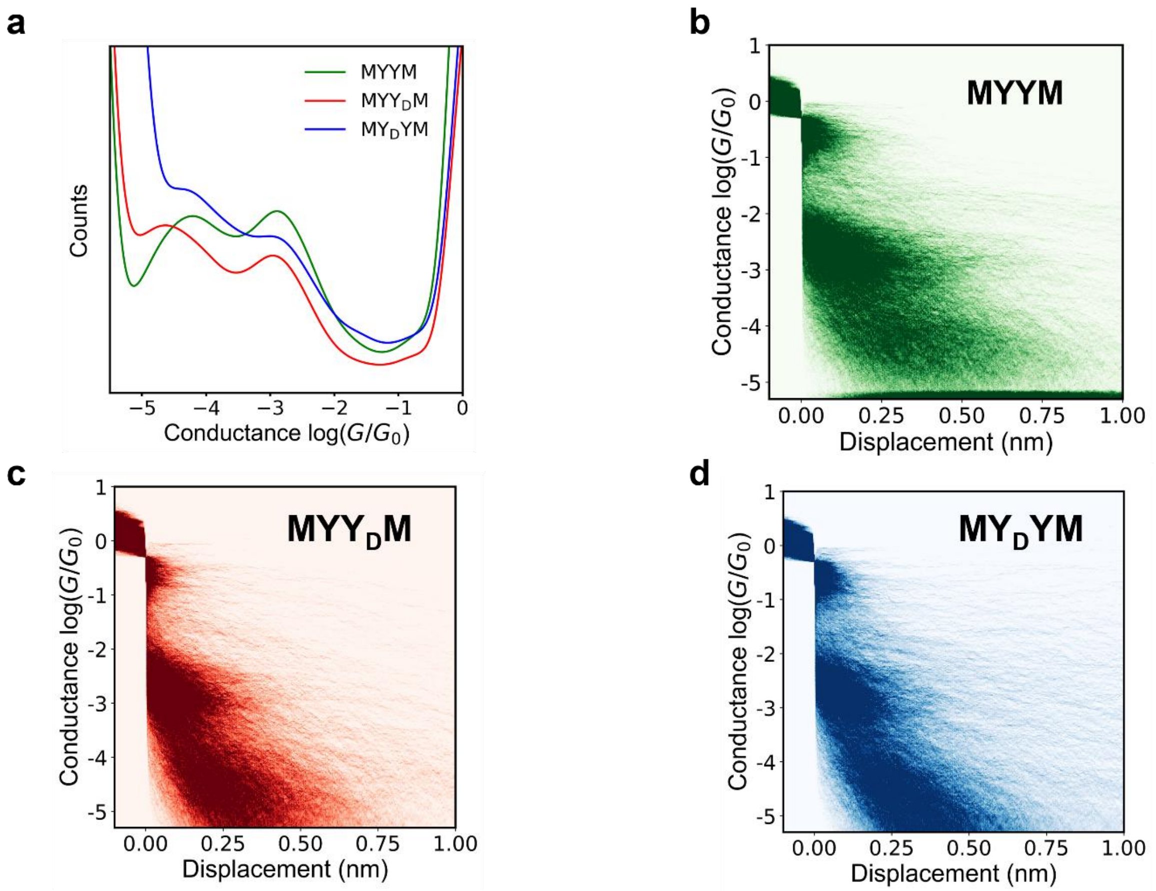

**Supplementary Figure 22**: Scanning tunneling microscope-break junction (STM-BJ) experiments for tyrosine-based tetrapeptides. (a) 1D conductance histograms for peptides MYYM, $MYY_DM$, and $MY_DYM$. (b), (c), (d) 2D conductance histograms for peptides MYYM, $MYY_DM$, and $MY_DYM$. Data were obtained using 0.1 mM concentrations of peptides in water at 250 mV applied bias across ensembles of at least 5000 single molecules.

**Supplementary Table S1**: High conductance and low conductance peaks for tyrosine-based tetrapeptides. The conductance value, in log scale, for each peak (high conductance and low conductance) is determined from the center position of a Lorentzian fit to the peak.

| **Sequence** | **High Conductance peak** [log($G/G_0$)] | **Low Conductance peak** [log($G/G_0$)] |
|---|---|---|
| MYYM | -2.80 | -4.32 |
| MYY$_D$M | -2.72 | -4.87 |
| MY$_D$YM | -2.75 | -4.49 |

**Supplementary Table S2**: High conductance and low conductance peaks for tyrosine-based pentapeptides. The conductance value, in log scale, for each peak (high conductance and low conductance) is determined from the center position of a Lorentzian fit to the peak.

| **Sequence** | **High Conductance peak** [log($G/G_0$)] | **Low Conductance peak** [log($G/G_0$)] |
|---|---|---|
| MYYYM | -2.92 | -4.04 |
| MYY$_D$YM | -2.80 | -4.80 |

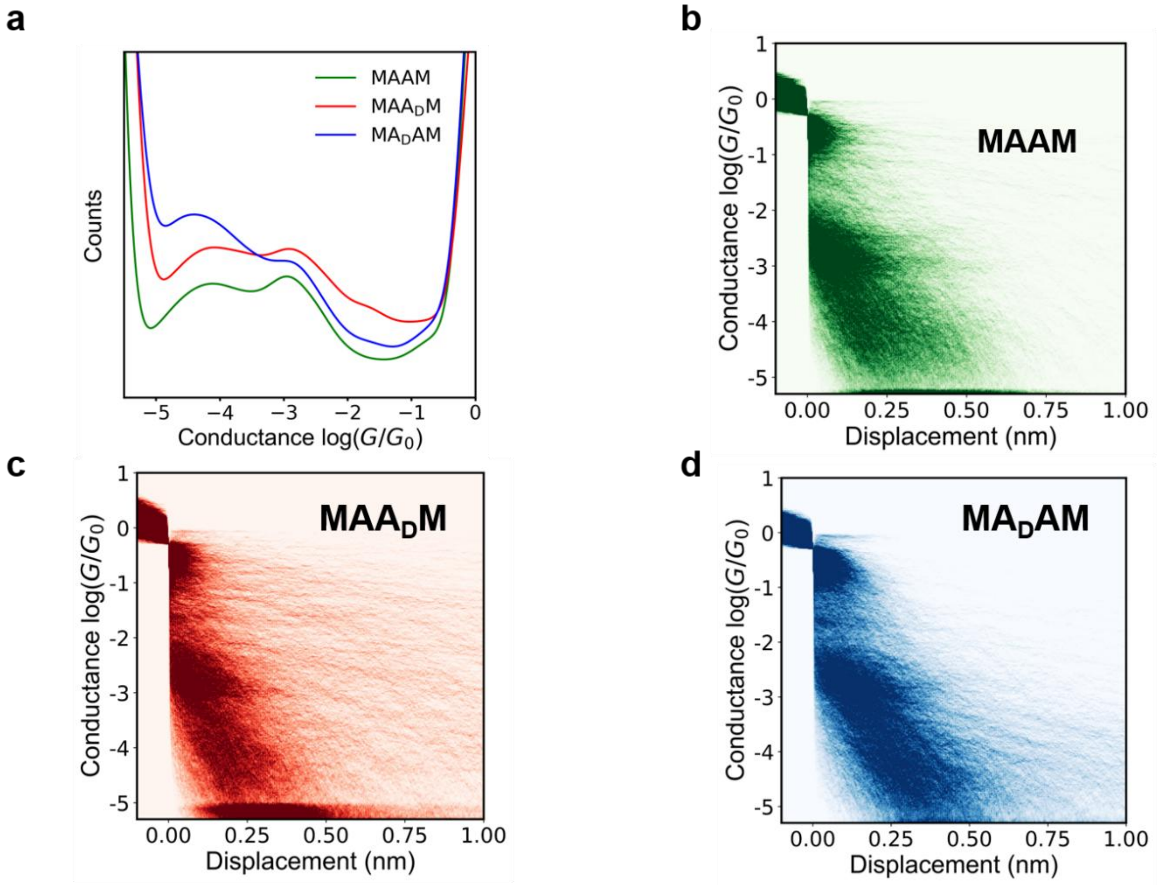

**Supplementary Figure 23**: Scanning tunneling microscope-break junction (STM-BJ) experiments for tyrosine-based tetrapeptides. (a) 1D conductance histograms for peptides MAAM, MAA$_D$M, and MA$_D$AM. (b), (c), (d) 2D conductance histograms for peptides MAAM, MAA$_D$M, and MA$_D$AM. Data were obtained using 0.1 mM concentrations of peptides in water at 250 mV applied bias across ensembles of at least 5000 single molecules.

**Supplementary Table S3**: High conductance and low conductance peaks for alanine-based tetrapeptides. The conductance value, in log scale, for each peak (high conductance and low conductance) is determined from the center position of a Lorentzian fit to the peak.

| **Sequence** | **High Conductance peak** [log(G/$G_0$)] | **Low Conductance peak** [log(G/$G_0$)] |
|---|---|---|
| MAAM | -2.86 | -4.22 |
| MAA$_D$M | -2.71 | -4.40 |
| MA$_D$AM | -2.75 | -4.27 |

**Supplementary Table S4**: High conductance and low conductance peaks for alanine-based pentapeptides. The conductance value, in log scale, for each peak (high conductance and low conductance) is determined from the center position of a Lorentzian fit to the peak.

| **Sequence** | **High Conductance peak** [log(G/$G_0$)] | **Low Conductance peak** [log(G/$G_0$)] |
|---|---|---|
| MAAAM | -2.91 | -4.63 |
| MAA$_D$AM | -2.83 | -4.88 |

### S7. Molecular dynamics (MD) simulations

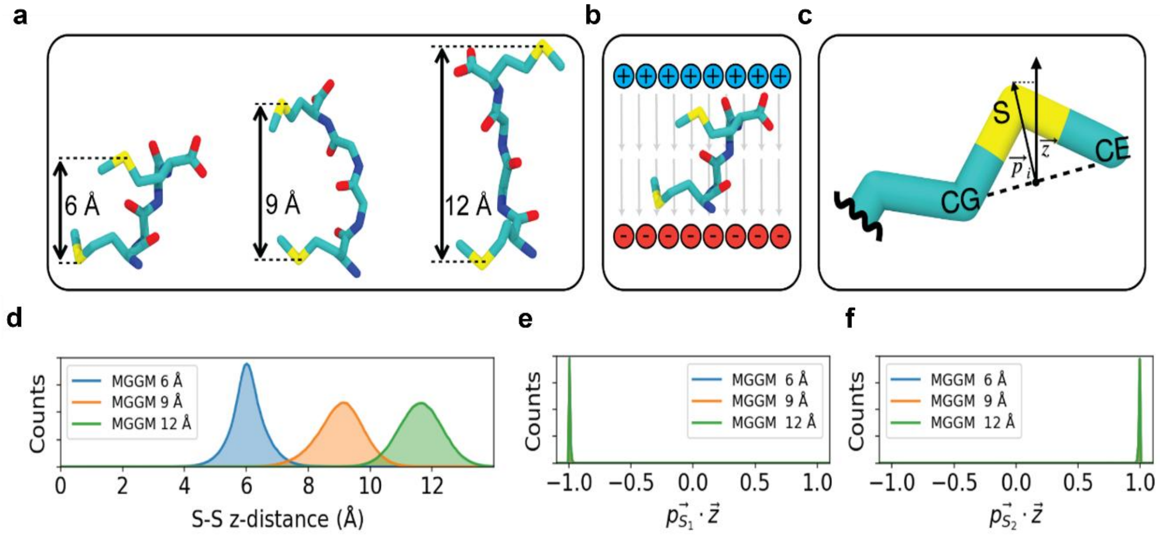

**Supplementary Figure 24:** Schematic illustration of implicit potentials utilized for MD simulations as described in our prior work. [4] (a), (d) Inter-anchor displacement potentials and resulting distributions at 6 Å, 9 Å, and 12 Å holding stages, corresponding to Equation 1 (main text). (b) Applied electric field defined in Equation 2 (main text). (c), (e), (f) Sulfur-orienting potential and resulting distributions defined in Equation 3 (main text).

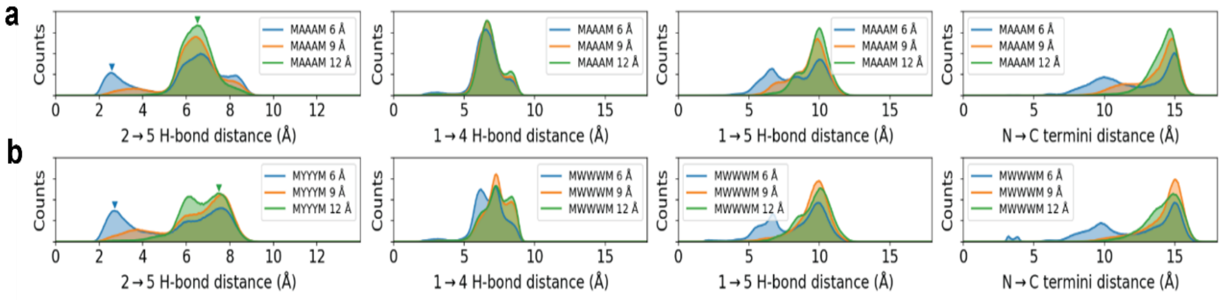

**Supplementary Figure 25:** MD simulation results for homochiral pentapeptides. Backbone hydrogen bonding (H-bonding) distance distributions for (a) MAAAM and (b) MYYYM, indicating elimination of intramolecular H-bonds at higher displacement holding stages. Figures from left to right indicate different backbone intramolecular H-bonds (2 → 5 backbone hydrogen bond, 1 → 4 backbone hydrogen bond, 1 → 5 backbone H-bond, and the distance between N → C termini. This methodology of utilizing MD simulations for interpreting backbone H-bonding interactions is based on prior work[4].

### S8. NEGF-DFT and quantum calculations

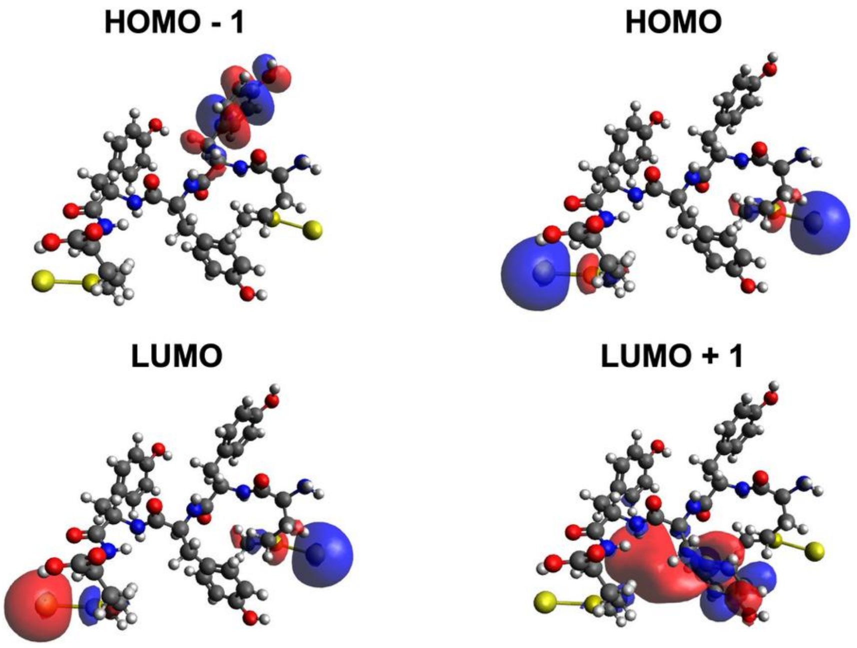

**Supplementary Figure 26:** Frontier molecular orbital analysis for MYYYM at a holding distance of 12 Å, corresponding to the low-conductance state or the primary structure-based electron transport pathway observed in single-molecule experiments.

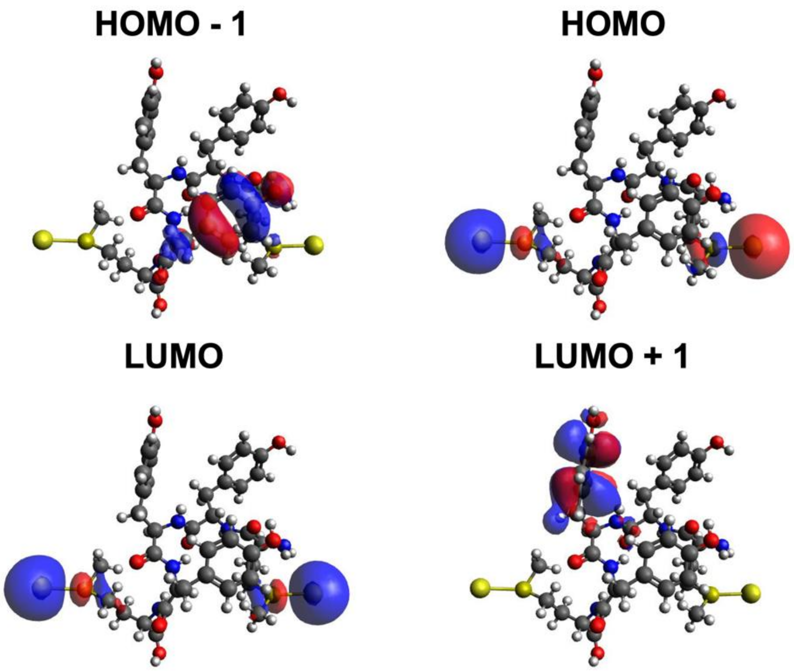

**Supplementary Figure 27:** Frontier molecular orbital analysis for $MYY_DYM$ at a holding distance of 12 Å, corresponding to the low-conductance state or the primary structure-based electron transport pathway observed in single-molecule experiments.

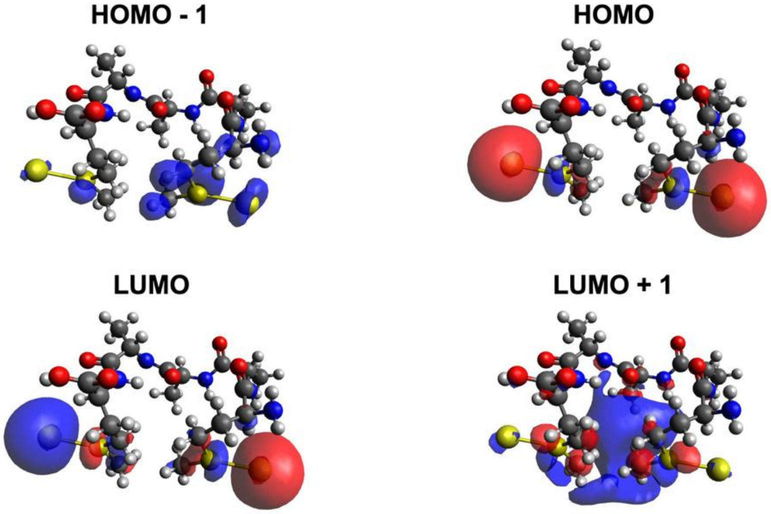

**Supplementary Figure 28:** Frontier molecular orbital analysis for MAAAM at a holding distance of 6 Å, corresponding to the high-conductance state or the secondary structure-based electron transport pathway observed in single-molecule experiments.

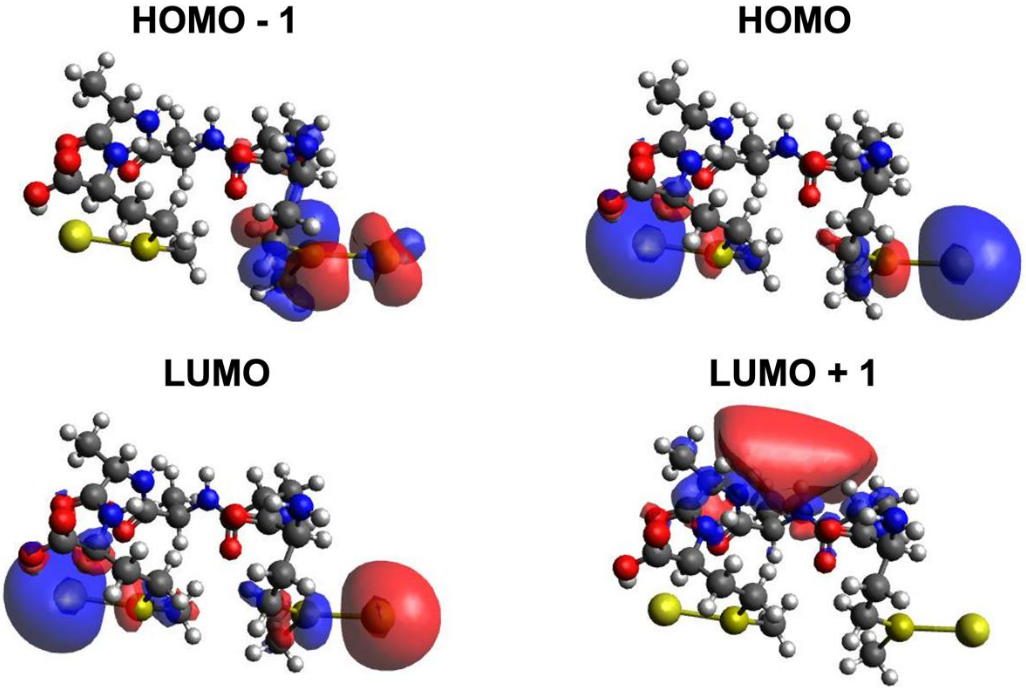

**Supplementary Figure 29:** Frontier molecular orbital analysis for MAA$_D$AM at a holding distance of 6 Å, corresponding to the high-conductance state or the secondary structure-based electron transport pathway observed in single-molecule experiments.

**Supplementary Table S5**: Frontier molecular orbital analysis for peptides. HOMO-1, HOMO, LUMO, and LUMO+1 values are reported for MAAAM, $MAA_DAM$, MYYYM, and $MYY_DYM$.

| Peptide Sequence | HOMO – 1 (eV) | HOMO (eV) | LUMO (eV) | LUMO + 1 (eV) |
|---|---|---|---|---|
| MAAAM | -9.2090 | -4.9776 | -3.5708 | 0.9629 |
| $MAA_DAM$ | -8.9030 | -4.5220 | -3.1577 | 0.8229 |
| MYYYM | -8.6820 | -4.7173 | 2.9862 | 7.5236 |
| $MYY_DYM$ | -8.5837 | -4.3587 | 3.3233 | 7.3348 |

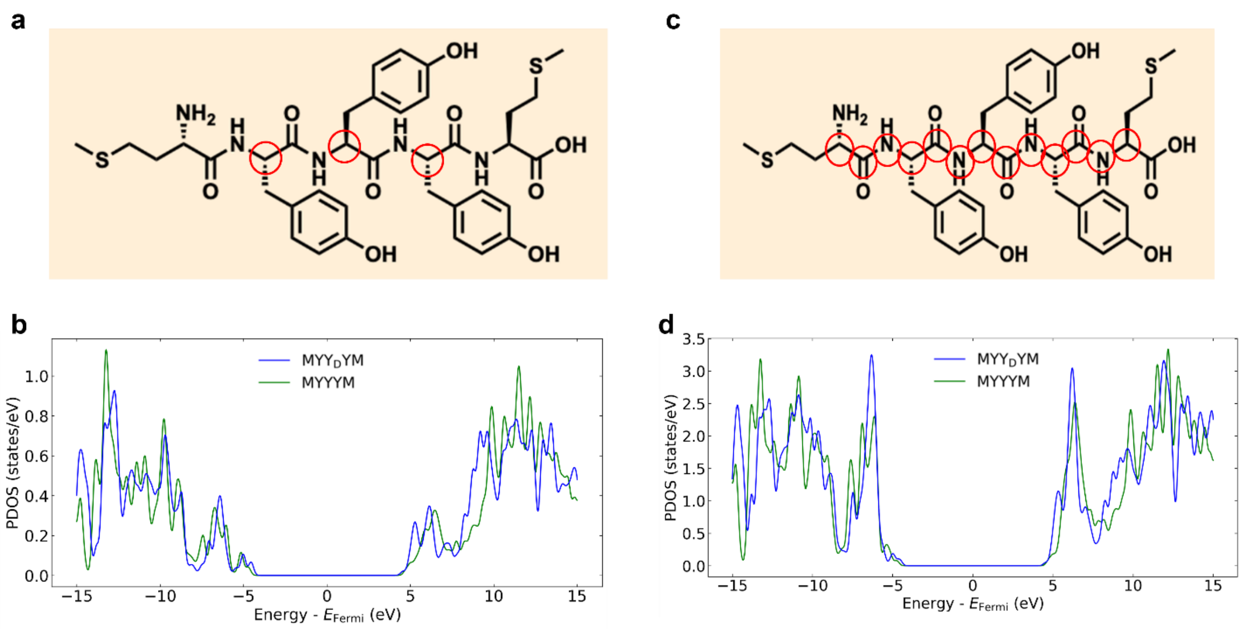

**Supplementary Figure 30:** Projected density of states (PDOS) calculations for MYYYM and MYY$_D$YM. (a), (b) Site specific PDOS calculations for carbon atoms on the peptide backbone for which the side chain is attached. (c), (d) PDOS for all backbone heavy atoms.

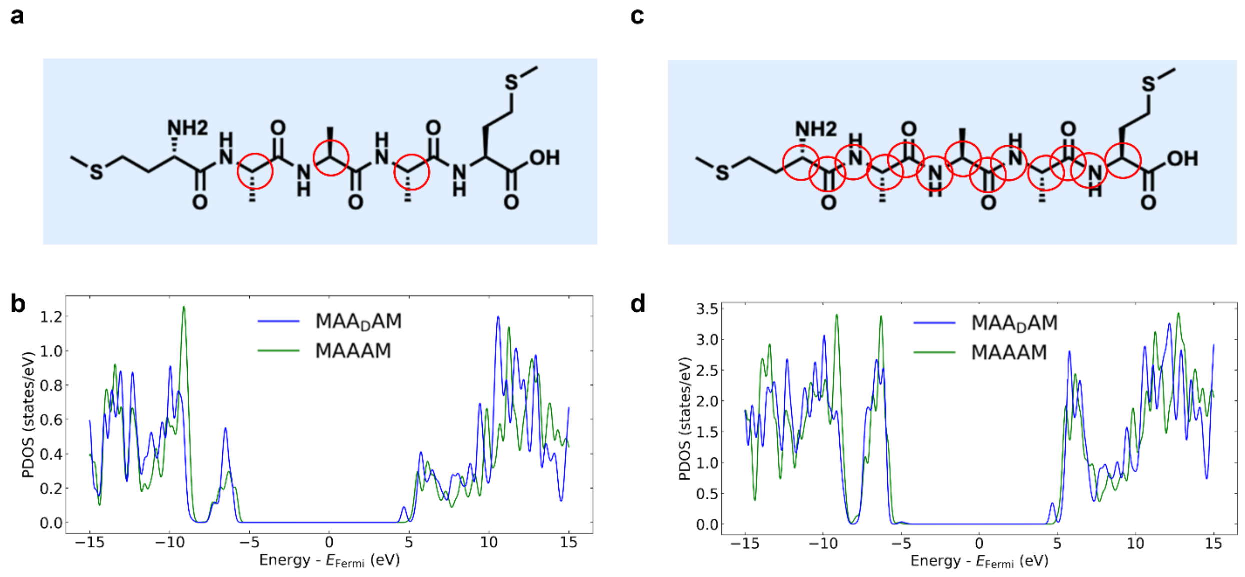

**Supplementary Figure 31:** Projected density of states (PDOS) calculations for MAAAM and MAA$_D$AM. (a), (b) Site specific PDOS calculations for carbon atoms on the peptide backbone for which the side chain is attached. (c), (d) PDOS for all backbone heavy atoms.

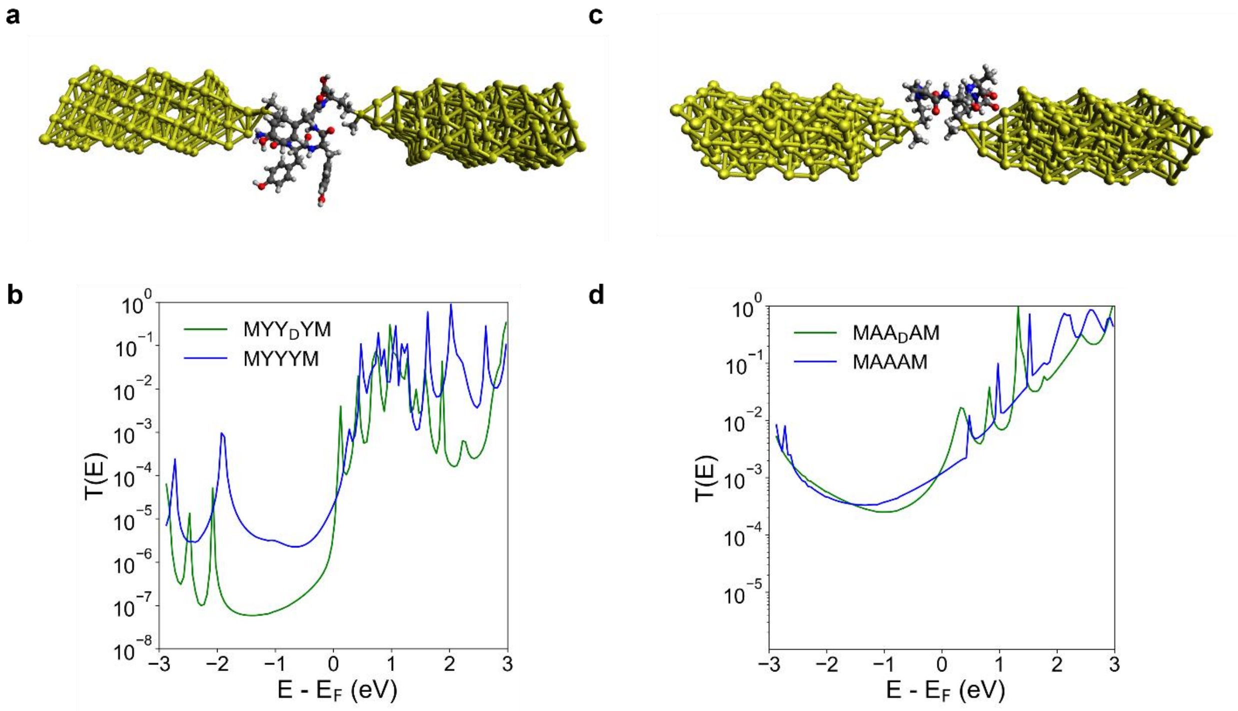

**Supplementary Figure 32:** NEGF-DFT calculations for MYYYM, MYY$_D$YM, MAAAM, and MAA$_D$AM. (a) Schematic of the MYY$_D$YM molecular junction with gold electrodes. (b) Electron transmission plots for MYYYM and MYY$_D$YM. (c) Schematic of the MAA$_D$AM molecular junction with gold electrodes. (d) Electron transmission plots for MAAAM and MAA$_D$AM. All NEGF-DFT calculations were performed using hollow gold electrodes in the absence of charges.

**Supplementary Table S6:** Computed electron transmission values from NEGF-DFT calculations and corresponding experimental transmission values obtained from STM-BJ measurements for selected peptide sequences for the low conductance state for the tyrosine-based peptides (12 Å end-to-end distance along the pulling axis) at the Fermi energy level. The NEGF-DFT calculations are performed using a hollow electrode geometry and without the presence of charges.

| Sequence | Computed electron transmission value (using NEGF-DFT) | Experimental electron transmission value (using STM-BJ experiments) |
|---|---|---|
| MYYYM | $2.1 \times 10^{-5}$ | $9.1 \times 10^{-5}$ |
| MYY$_D$YM | $5.7 \times 10^{-6}$ | $1.6 \times 10^{-5}$ |

**Supplementary Table S7:** Computed electron transmission values from NEGF-DFT calculations and corresponding experimental transmission values obtained from STM-BJ measurements for selected peptide sequences for the low conductance state for the alanine-based peptides (6 Å end-to-end distance along the pulling axis) at the Fermi energy level. The NEGF-DFT calculations are performed using a hollow electrode geometry and without the presence of charges.

| Sequence | Computed electron transmission value (using NEGF-DFT) | Experimental electron transmission value (using STM-BJ experiments) |
|---|---|---|
| MAAAM | $1.2 \times 10^{-3}$ | $1.2 \times 10^{-3}$ |
| MAA$_D$AM | $1.5 \times 10^{-3}$ | $1.4 \times 10^{-3}$ |

### S9. References

1. Li, S.; Yu, H.; Schwieter, K.; Chen, K.; Li, B.; Liu, Y.; Moore, J. S.; Schroeder, C. M. Charge transport and quantum interference effects in oxazole-terminated conjugated oligomers. *Journal of the American Chemical Society* **2019**, 141 (40), 16079–16084.
2. Li, B.; Yu, H.; Montoto, E. C.; Liu, Y.; Li, S.; Schwieter, K.; Rodríguez-López, J.; Moore, J. S.; Schroeder, C. M. Intrachain charge transport through conjugated donor–acceptor oligomers. *ACS Applied Electronic Materials* **2018**, 1 (1), 7–12.
3. Venkataraman, L.; Klare, J. E.; Tam, I. W.; Nuckolls, C.; Hybertsen, M. S.; Steigerwald, M. L. Single-molecule circuits with well-defined molecular conductance. *Nano Letters* **2006**, 6 (3), 458–462.
4. Samajdar, R.; Meigooni, M.; Yang, H.; Li, J.; Liu, X.; Jackson, N. E.; Mosquera, M. A.; Tajkhorshid, E.; Schroeder, C. M. Secondary structure determines electron transport in peptides. *Proceedings of the National Academy of Sciences* **2024**, 121 (32), e2403324121.
5. Liu, Xiaolin; Yang, Hao; Harb, Hassan; Samajdar, Rajarshi; Woods, Toby J.; Lin, Oliver; Chen, Qian; Romo, Adolfo I. B.; Rodríguez-López, Joaquín; Assary, Rajeev S.; Moore, Jeffrey S.; Schroeder, Charles M. Shape-persistent ladder molecules exhibit nanogap-independent conductance in single-molecule junctions. *Nature Chemistry* **2024**, 16, 1-9.
6. Samajdar, R.; Yang, H.; Yi, S.; Wang, C.-I.; Putnam, S. T.; Pence, M. A.; Lindsay, G. S.; Meigooni, M.; Liu, X.; Ren, J.; Moore, J. S.; Tajkhorshid, E.; Gewirth, A. A.; Rodríguez-López, J.; Jackson, N. E.; Schroeder, C. M. Electrochemically mediated Au–C($sp^2$) anchors for molecular electronics. *The Journal of Physical Chemistry C* **2025**, 129 (39), 17458–17471.
7. Nagahara, L. A.; Thundat, T.; Lindsay, S. M. Preparation and characterization of STM tips for electrochemical studies. *Review of Scientific Instruments* **1989**, 60 (10), 3128–3130.
8. Tien, M. Z., Sydykova, D. K., Meyer, A. G., & Wilke, C. O. PeptideBuilder: A simple Python library to generate model peptides. *PeerJ* **1**, e80 (2013).
9. Humphrey, W., Dalke, A., & Schulten, K. VMD: visual molecular dynamics. *J. mol. graphics* **14**, 33-38 (1996).
10. Brooks, B. R.; Brooks, C. L., III; Mackerell, A. D., Jr; Nilsson, L.; Petrella, R. J.; Roux, B.; Won, Y.; Archontis, G.; Bartels, C.; Boresch, S.; Caflisch, A. CHARMM: the biomolecular simulation program. *Journal of Computational Chemistry* **2009**, 30 (10), 1545–1614.
11. Huang, J.; Rauscher, S.; Nawrocki, G.; Ran, T.; Feig, M.; De Groot, B. L.; Grubmüller, H.; MacKerell, A. D., Jr. CHARMM36m: an improved force field for folded and intrinsically disordered proteins. *Nature Methods* **2017**, 14 (1), 71–73.
12. Eastman, P.; Swails, J.; Chodera, J. D.; McGibbon, R. T.; Zhao, Y.; Beauchamp, K. A.; Wang, L.-P.; Simmonett, A. C.; Harrigan, M. P.; Stern, C. D.; Wiewiora, R. P.; Brooks, B. R.; Pande, V. S. OpenMM 7: rapid development of high performance algorithms for molecular dynamics. *PLoS Computational Biology* **2017**, 13, e1005659.

13. Zhang, Z.; Liu, X.; Yan, K.; Tuckerman, M. E.; Liu, J. Unified efficient thermostat scheme for the canonical ensemble with holonomic or isokinetic constraints via molecular dynamics. *Journal of Physical Chemistry A* **2019**, 123, 6056–6079.
14. Darden, Tom.; Darrin York.; Lee Pedersen. "Particle mesh Ewald: An N log (N) method for Ewald sums in large systems." *Journal of Chemical Physics* **1993**, 98, 10089–10092.
15. Shaw, D. E.; Deneroff, M. M.; Dror, R. O.; Kuskin, J. S.; Larson, R. H.; Salmon, J. K.; Young, C.; Batson, B.; Bowers, K. J.; Chao, J. C.; Eastwood, M. P. Anton: A special-purpose machine for molecular dynamics simulation. *Communications of the ACM* **2008**, 51, 91–97.
16. Shirts, M.; Pande, V. S. Screen savers of the world unite! *Science* **2000**, 290, 1903–1904.
17. Brandbyge, M.; Mozos, J.-L.; Ordejón, P.; Taylor, J.; Stokbro, K. Density-Functional Method for Nonequilibrium Electron Transport. *Phys. Rev. B* **2002**, *65* (16), 165401.
18. Soler, J. M.; Artacho, E.; Gale, J. D.; García, A.; Junquera, J.; Ordejón, P.; Sánchez-Portal, D. The SIESTA Method for Ab Initio Order-N Materials Simulation. *J. Phys.: Condens. Matter* **2002**, *14* (11), 2745–2779.
19. Papior, N.; Lorente, N.; Frederiksen, T.; García, A.; Brandbyge, M. Improvements on Non-Equilibrium and Transport Green Function Techniques: The Next-Generation TranSIESTA. *Comput. Phys. Commun.* **2017**, *212*, 8–24.
20. Perdew, J. P.; Burke, K.; Ernzerhof, M. Generalized Gradient Approximation Made Simple. *Phys. Rev. Lett.* **1996**, *77* (18), 3865–3868. DOI: 10.1103/PhysRevLett.77.3865.
21. Sharma, B.; Asher, S. A. UV resonance Raman investigation of the conformations and lowest energy allowed electronic excited states of tri- and tetraalanine: Charge transfer transitions. *The Journal of Physical Chemistry B* **2010**, 114, 6661–6668.
22. Heller, A.; Rönitz, O.; Barkleit, A.; Bernhard, G.; Ackermann, J.-U. Complexation of europium (III) with the zwitterionic form of amino acids studied with ultraviolet–visible and time-resolved laser-induced fluorescence spectroscopy. *Applied Spectroscopy* **2010**, 64, 930–935.
